# Assembly and symmetry of the fungal E3BP-containing core of the Pyruvate Dehydrogenase Complex

**DOI:** 10.1101/2020.08.13.232140

**Authors:** B. O. Forsberg, S. Aibara, R. J. Howard, N. Mortezaei, E. Lindahl

**Affiliations:** Department of Biochemistry and Biophysics, Science for Life Laboratory, Stockholm University, 17165 Solna, Sweden; Department of Molecular Biology, Max Planck Institute for Biophysical Chemistry, 37077 Göttingen, Germany; Vironova AB, 11330 Stockholm, Sweden; Department of Applied Physics, Swedish eScience Research Center, KTH Royal Institute of Technology, 17168 Solna, Sweden

## Abstract

The pyruvate dehydrogenase complex (PDC) is a central component of all aerobic respiration, connecting glycolysis to mitochondrial oxidation of pyruvate. Despite its central metabolic role, its precise composition and means of regulation remain unknown. To explain the variation in stoichiometry reported for the E3-recruiting protein X (PX) in the fungal PDC, we established cryo-EM reconstructions of the native and recombinant PDC from the filamentous fungus and model organism *Neurospora crassa*. We find that the PX C-terminal domain localizes interior to the E2 core. Critically, we show that two distinct arrangements of a trimeric oligomer exists, which both result in strict tetrahedral symmetry of the PDC core interior. Both oligomerization and volume occlusion of the PDC interior by PX appears to limit its binding stoichiometry, which explains the variety of stoichiometries found previously for *S. cerevisiae*. This also suggests that the PX oligomer stability and size are potential mechanisms to dynamically adjust PDC compostion in response to external cues. Moreover, we find that the site where PX binds is conserved within fungi but not mammals, suggesting that it could be therapeutically targeted. To this end, we also show that a PX knockout results in loss of activity through dysfunctional E3 recruitment, leading to severely impaired *N. crassa* growth on sucrose. The fungal PDC is thus shown to be fundamentally similar to the mammalian PDC in function but subject to other conditions of possible regulation, conditioned by a steric restrictions imposed by the symmetry of the PDC and its components.

---

Anaerobic glycolysis catabolizes glucose through a series of electron transfer reactions that produce adenosine triphosphate (ATP) and the terminal electron acceptor pyruvate. The pyruvate dehydrogenase complex (PDC) catalyzes the further formation of acetyl-coenzyme A (acetyl-CoA) from pyruvate, enabling the tricarboxylic acid (TCA) cycle that sustains oxidative phosphorylation. Consequently, deficiency or malfunction of the PDC profoundly impacts metabolic fitness and normal development^1^, and is also associated with severe metabolic disorders such as Leigh syndrome and episodic ataxia^2^. In cancer biology, the PDC attracts attention because aerobic glycolysis is up-regulated in many cancers, while inhibition of the PDC results in decreased mitochondrial glucose oxidation, known as the Warburg effect^3,4^. Decreased mitochondrial activity and reduction of reactive oxygen species (ROS) generated by the PDC^5^ both inhibit fundamental apoptotic pathways^6^, whereas activation of the PDC in symptomatic melanomas has in fact been shown to restore mitochondrial activity and induced senescence^7^.

The PDC is a multienzyme complex consisting of three components: E1, E2 and E3. E1 catalyzes the decarboxylation of pyruvate and transfers the resultant acetyl-group to an E2 carrier domain. E2 then covalently links this acetyl group to CoA through its catalytic domain. Finally, E3 catalyzes the re-oxidation of the E2 carrier domain lipoyl moeity, allowing the chain of reactions to repeat. Without exception, the PDC is organised by non-covalent recruitment of E1 and E3 to a large core built from E2 catalytic domains. The tethering of E1 and E3 to the E2 core by disordered linking regions produces a flexible complex that is challenging to reconstruct faithfully, since any such reconstruction is a convolution of the true object with the probability distribution describing the relative positions of its components (among other factors). A component with a well-defined position will result in an interpretable reconstructed convolution dominated by the true component structure, regardless of nominal resolution. In contrast, more flexible components carry the probability distribution as a more dominant characteristic and are not interpretable as faithful yet low-pass filtered reconstructions, as previously assumed^8–10^. A global view of fully assembled PDC, or comprehensive knowledge of its flexibility and transient contacts has never been presented. Nevertheless, structures of isolated PDC components^11,12^, and core assemblies with high symmetry^13–15^ have been determined.

The co-localization of the participating enzymes and substrates increases the overall rate by minimizing diffusion, and tightly couples the chain of reactions. The tight coupling in turn enables rapid and reversible regulation of the reaction cycle through phosphorylation of E1 by kinases and phosphatases that are weakly bound to the PDC^12^, adapting its activity to current metabolic requirements^16,17^. Finer control over metabolic flux is also afforded by adjusting the relative proportions of the PDC components, regulating its overall activity. The relative proportions of E1 and E3 follow their respective affinity for the recruitment domains of the core. In mammals this is further exploited through the expression of a catalytically inactive E2 paralog called E3-binding protein (E3BP) that partially substitutes E2 and selectively recruits E3^18^. The fungal PDC also expresses an E2-homologous protein to recruit E3^19,20^, originally known as “protein X” or the “X component” (PX). There is however no obvious similarity in the C-terminal domains between mammalian E3BP and fungal PX. This suggests that a reduction in E2 activity by replacement with catalytically inactive subunits is unlikely in fungal PDC, and decoupled from E3 recruitment. It has not yet been possible to show how these E3-specific components assemble in the PDC core for any mammalian or fungal species, but models and stoichiometric measurements have suggested E2:PX stoichiometries of 2:1^21^, 4:1^22,23^, and 5:1^24^. The challenge to determine how this regulation is dictated by structure, lies in the suppression of relatively small asymmetric features by a largely symmetric complex, here termed *implicit symmetrization*.

To show how fungal PX is incorporated into the core assembly and stoichiometrically regulated, we present a structural analysis of the PDC from the fungal model organism *Neurospora crassa*. We consider the implicit symmetrization caused by the icosahedral E2 core and are able to determine tetrahedral reconstructions, showing how the fungal PDC departs from the dominating symmetry. The tetrahedral-symmetric arrangement of PX interior to the icosahedral PDC core dictates a complex composition similar to that suggested for mammalian PDC. Based on its oligomeric state and size, we also identify factors that offer a simple explanation for the range of previously presented PDC component ratios in fungi. We confirm that PX is critical to *N. crassa* pyruvate metabolism and that by extension the stoichiometric ratio of the complex components pivotal to overall activity. Any change in PX oligomer stability and/or size would therefore affect PDC component stoichiometry, which could be utilized to allow fungi to tune PDC component activity in a dynamic fashion. Further, we discuss the fidelity of the established classification and exemplify how marginalized reconstruction leads to uncertainty in the classified proportions of cryo-EM particles, which is crucial to the interpretation of the present results.

## 1 Results

### 1.1 Structure determination of a non-icosahedral N. crassa PDC core component

In order to characterize the fungal PDC, we purified intact complexes from *N. crassa* mitochondria and reconstructed its native structure from single-particle cryo-electron microscopy data. This showed a dodecahedron-shaped core with icosahedral symmetry, composed of 60 E2 core-forming domains decorated with many flexibly tethered enzymes surrounding it (Fig. 1A). To improve the reconstruction, icosahedral symmetry was enforced. This reconstruction allowed identification of E2 residues 232-458, including part of the N-terminal linker that extends towards the recruitment domain (peripheral subunit-binding domain, PSBD). Interior to the core assembly, we found an apparently ordered, non-icosahedral PDC component (Fig. 1A), which appears consistent with a mixture of states, incorrectly averaged in the icosahedral asymmetric unit (I-ASU). In order to reconstruct this component faithfully, we attempted to avoid over-symmetrization through use of a lower symmetry group. While this alone did not improve the interpretability of the interior density for any icosahedral sub-symmetry, 3D classification coupled with the application of tetrahedral symmetry yielded a correctly averaged ASU, evidenced by a protein-like density (Fig. 1D) and consistent background in the complementary interior volume. A preliminary analysis of the interior assembly suggested that an alternate tetrahedral configuration should exist with equal probability. Through repeated classification using only the core assembly as an initial reference, such a state was indeed observed. Three distinct classes of the T-ASU were consistently observed, partially reconciling the over-symmetrized appearance of the internal density within the I-ASU (Fig. 1B). All such classes show an identical E2 core scaffold with the same handedness, differing only in their interior content. The first class has four interior, threefold symmetric densities, each immediately interior to an E2 trimer. Each such threefold interior density appears as a basket which hangs underneath (interior to) an associated E2 trimer. They are arranged such that they associate with E2 trimers along positive direction of each threefold axis of the tetrahedral symmetry (Fig. 1C). The second class shows interior densities identical to the first, but arranged with E2 trimers along the negative direction of each threefold symmetry axis, resulting in a distinct relative arrangement that cannot be equated to the first by an affine transformation. These classes are therefore denoted tetrahedral classes X and Y, respectively. The third class appears to represent an over-symmetrized T-ASU (Fig. 1), similar to the I-ASU, which we term the non-tetrahedral class N. To approach a quantification of the distribution of data across these classes, a three-class classification was conducted using the above reconstructions as simultaneous and separate input references. This assigned approximately 30% of the input particles to either of the classes X and Y, and the remaining 40% to class N.

**Figure 1:**
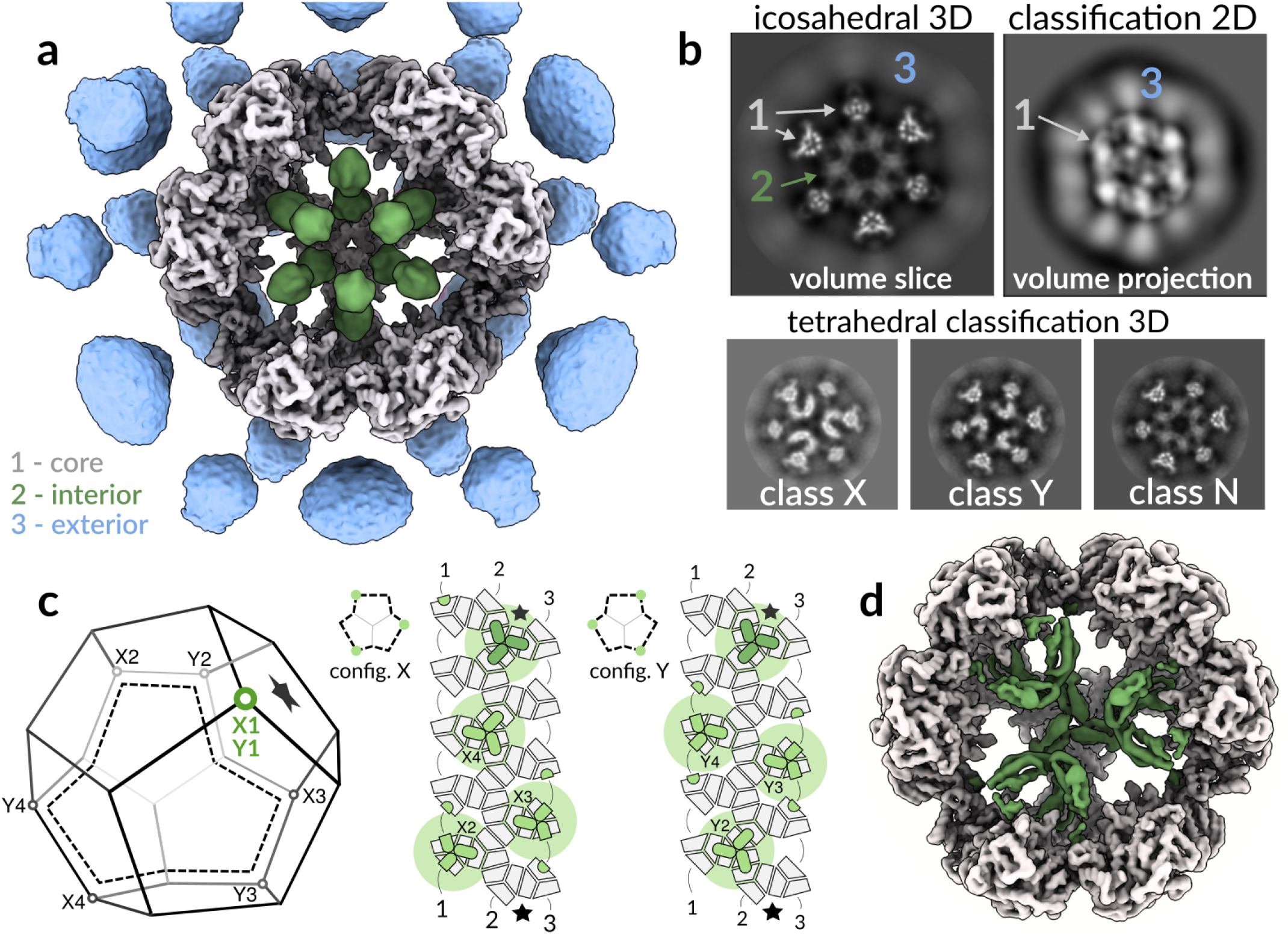
The *N. crassa* PDC has a core-internal protein with tetrahedral optimal packing. **a**, Reconstruction of the intact native PDC icosahedral Asymmetric unit (I-ASU) shows symmetrization artifacts, both exterior 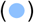 and interior 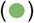 to the core 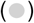, predominantly along five-fold (interior and exterior) and twofold (exterior) symmetry axes. **b**, Whereas the exterior artifact appears to represent a continuous degree of freedom resulting in a blurred but elevated density, the interior component is consistent with a discrete mixture of states. Classification of the tetrahedral ASU (T-ASU) resolves this mixture into three classes of interest: tetrahedral classes X and Y, and class N that still appears to be over-symmetrized. **c**, A schematic dodecahedron (left) illustrates which four E2 trimers (vertices) are occupied by interior density in each of classes X and Y. These arrangements are also shown mapped to an unfolded view of the E2 core (right), where each E2 monomer is shown in grey, and each interior monomer is shown in green. These monomers form threefold symmetric oligomers. The effective volumetric occlusion of the core interior is schematically illustrated by a background circle - two interior components cannot be placed closer than shown here. **d**, A cut-away view illustrates the overall reconstruction of the tetrahedral asymmetric unit (T-ASU) using class Y, and the protein-like density (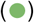) interior to the PDC core.

To explain the prevalence of the tetrahedral arrangement in native PDC, we next evaluated what steric restrictions are associated with the interior density volume occlusion. We computationally isolated a single threefold interior density, and examined which configurations of multiple such densities result in clashes. We find that no two E2-trimers separated by fewer than two intermediate-E2 trimers withinthe core scaffold can accommodate a structured interior basket density (Fig. S1D). Under this restriction it is possible to arrange four such interior densities in two unique arrangements (Fig. 1C, config. X and Y). These coincide with tetrahedral symmetry and correspond to each of the classes X and Y found by applying tetrahedral symmetry during cryo-EM classification. Note that we use the nomenclature *class* to describe a data subset and *configuration* for an idealized arrangement of interior density.

To further examine class N, that represents a seemingly non-tetrahedral class, we considered how the tetrahedral symmetry of the interior might break. First, it is possible to arrange three or even two interior trimers such that further additions can be made without incurring clashes (Fig. S1E, config 3S and 2S). We term these configurations *sub-optimal*. Second, if one or more available locations are unoccupied the interior would be *unsaturated* (e.g. Fig. S1E, config Y3U). Third, dimeric or monomeric interior densities are possible. Finally, a combination of any of the above cannot be ruled out. Attempts to re-classify the 40% of particles assigned to the non-tetrahedral class N under lower symmetry did not yield consistent results, but resulted in the reappearance of one or both of the tetrahedral classes when permitted by the enforced symmetry (D2,C3,C2,C1). In other symmetries (D5,D3,C5), the core is faithfully represented but the interior component appears over-symmetrized. We took this to indicate that class N contains particles with an unsaturated but optimally arranged interior, that tends to dominate re-classification and manifest as tetrahedral-like through explicit and/or implicit symmetrization. Any additional conflicting particles may be hidden by broad backprojection (see Discussion). No class of particles displaying sub-optimal saturation or asymmetric baskets was confidently identified, perhaps because the complete asymmetry of any such configuration would be out-competed by implicit symmetrization in the presence of the E2 core. Taken together, these observations indicate that the interior component prefers a threefold symmetric state, arranged according to an optimal geometry and near-saturation under the current (native) conditions.

We next refined the T-ASU of classes X and Y individually. The resolution of the interior density varies between 4.0-6.5 Å as assessed by local Fourier Shell Correlation (FSC, Fig. S2). Following local resolution filtering we confirm that classes X and Y both depict the same threefold-symmetric oligomer interior to the core. The only observable points of contact with the E2 core are in the twofold interface bridges connecting the density-associated E2 trimer to each of its three neighbors (Fig. 2A). E2 here provides a predominantly hydrophobic pocket, composed of the C-terminal residues that form the bridge and helix 2 (H2_E2_) (Fig. 2D). Notably, while H2_E2_ does not appear to be conserved among metazoa, its conservation in ascomycota indicates an essential function (Fig. 2D-E). Here, H2_E2_ contains a KLLK motif (K263–K266) with basic lysine residues oriented toward the interior density, although precise contacts remain to be identified. Additional conserved residues, including R268 and N272, interact with the substrate CoA^25^ rather than contributing to the interior-density interface, and should therefore not be considered constituents of the binding pocket.

**Figure 2:**
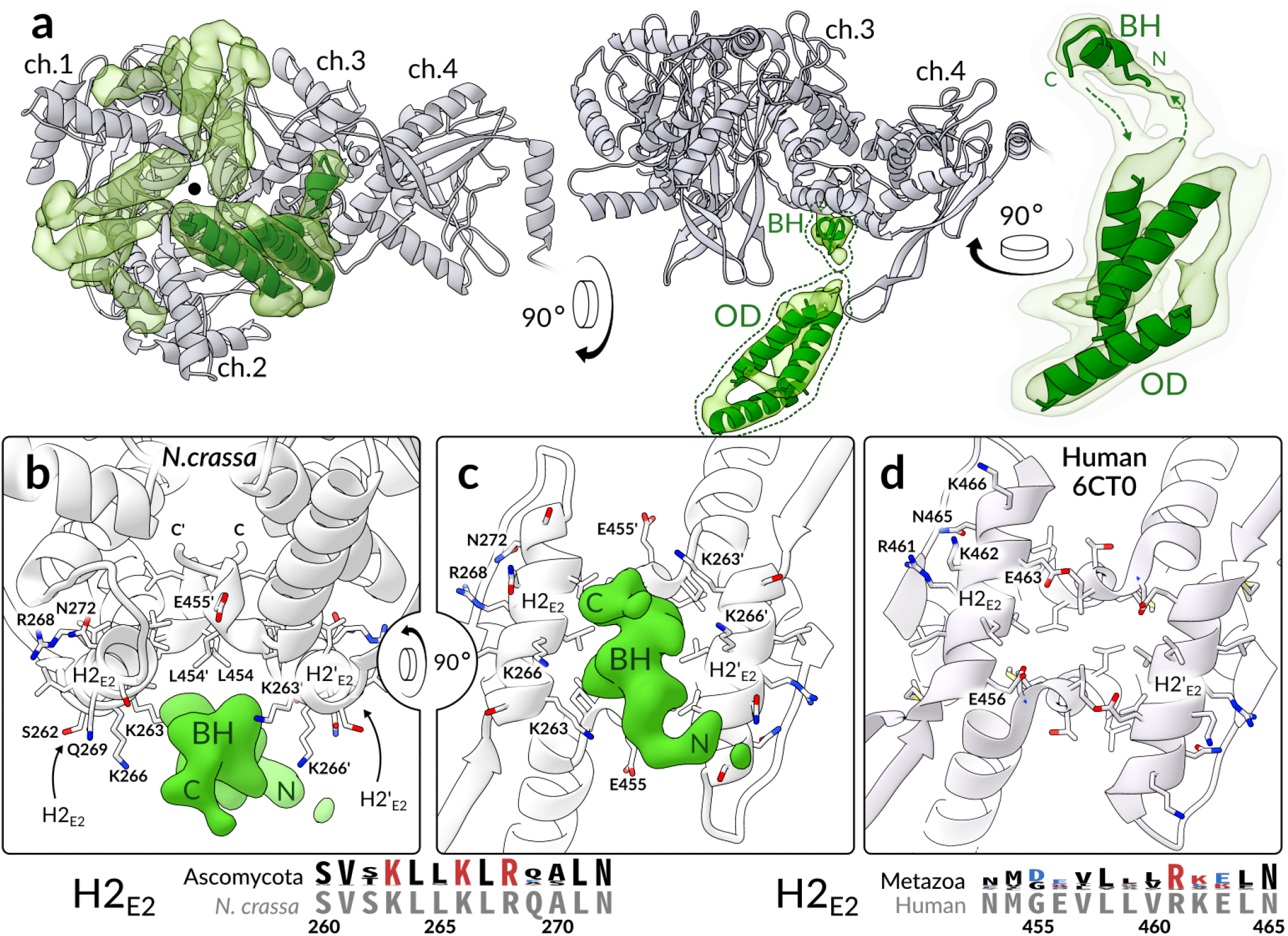
The E2-pair interface constitutes a conserved basic/hydrophobic binding pocket. **a**, E2 chains (gray 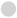) 1-3 form a threefold symmetric oligomer, and each of them additionally forms a twofold symmetric bridge to an adjacent E2 monomer (here labeled chain 4). A pocket inside each of these three twofold E2 bridges accommodates one binding helix (BH) of the threefold interior density (transparent green 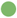). The bulk of the interior density forms the oligomer domain (OD) that extends N-terminally from the BH. The OD is consistent with a three-helix bundle, although secondary structure and connectivity remain ambiguous. Helices of approximately 20 residues are docked to illustrate this (green cartoon 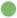). **b**, The BH binds asymmetrically to the symmetric pocket formed by the E2-bridge (Fig. S3). **c**, A rotated view shows that in ascomycota, pocket-lining lysines in H2_E2_ are conserved (sequence logo). **d**, In contrast, the human E2 interface is less basic and conserved, apart from residues involved in substrate interaction.

The segment of the interior density that binds to the E2-bridge pocket comprises a small (*∼*7 residues) alpha-helical segment, here referred to as the binding helix (BH). The backbone and direction of the BH could be traced from our reconstructions (Fig. 2B), but the resolution did not permit assignment of its primary sequence. The BH connects via its N-terminal end to the body of the threefold interior oligomer domain (OD), and via a weaker density to its C-terminus (Fig. 2A). Owing to its low reconstructed resolution, the precise fold of the OD is unclear, but it appears to be most consistent with a bundle of three alpha-helices of approximately 20 residues each (Fig. 2). The relatively low resolution of the OD density suggests it has some flexibility with respect to the E2 core. While this flexibility is expected given the delicate connection to the BH, the oligomer evidently supplies a sufficiently rigid or persistent steric obstacle to impose a tetrahedral symmetry on a large portion of the native PDC particles.

### 1.2 The core-interior density is a partial C-terminal domain of protein X

We next sought to confidently identify the protein occupying the PDC core interior. Mass spectrometry detected E1*α*, E1*β*, E2, E3, and Protein X, as well as kinases and phosphatases known to regulate the PDC (Fig. S4). E1 and E3 were excluded from consideration based on homologous structures being known and incompatible. To exclude both the lipoyl domain (LD) and PSBD of E2, we recombinantly expressed and purified *N. crassa* E2, and obtained a single-particle cryo-EM reconstruction. In contrast to the endogenous PDC, the interior of the recombinant E2 assemblies showed no evidence of a structured density subject to over-symmetrization (see Fig. 3A, S5) in the I-ASU. The interior did display a higher overall density compared to pure solvent, consistent with a flexible component, similar to the PDC exterior. We attribute this to the N-terminal PSBD and LD being free to diffuse into the core interior.

**Figure 3:**
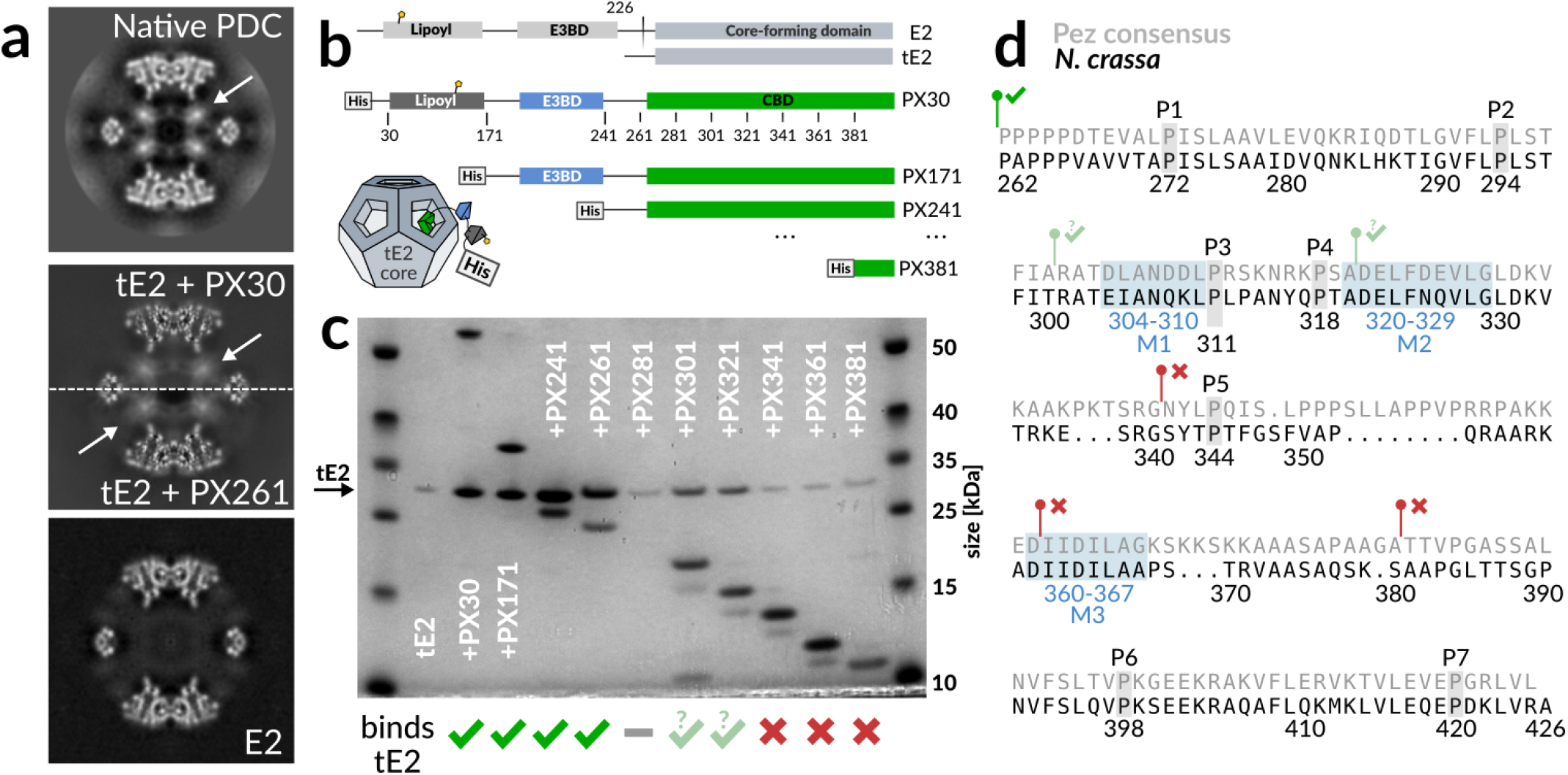
*N. crassa* PX is a monomer that oligomerizes upon binding to the PDC core. **a**, Comparison of over-symmetrized interior densities. The native PDC (top) was the only dataset wherein a tetrahedral arrangement of interior densities could be classified confidently. It is however clear that both a reconstituted tE2+PX30 as well as a co-expressed tE2+PX261 subcomplex (middle) display similar interior density-signatures under icosahedral (over-)symmetrization. The E2 core alone (bottom) does not show this signature. **b**, Schematic of recombinantly co-expressed tE2 and PX constructs used for binding assays. PX<N> indicates truncation of residues 1-N. **c**, SDS-PAGE showing co-purification of E2-core assemblies by His-tag affinity of PX constructs, illustrating a disproportionate decrease in bound E2 upon truncation past residue 280. PX281 exhibited a complete loss of expression or rapid degradation. Further truncation restored PX expression, but at reduced affinity or increased ratio. Binding was completely abolished (comparable to lone tE2, lane 2) following N-terminal truncation past residue 340. **d**, PX CBD sequence from *N. crassa* aligned to the the Pezizomycotina (Pez) consensus sequence. The seven conserved prolines are indicated, as well as the three conserved H2_E2_-complementary motifs M1-3, found in Pez. Notably for *N. crassa*, M1 does not agree well with the consensus motif (cf. Fig. S7). Green and red markers indicate the starting positions and binding of PX-constructs corresponding to panel **c**.

It seemed plausible that the C-terminal domain of fungal PX, which is homologous within fungi but not to the metazoan E3BP C-terminal domain, could account for the interior density. We therefore carried out separate recombinant expression of *N. crassa* E2 core-forming domain (tE2) and His-tagged PX variants, which could be co-purified by Ni-NTA affinity following brief incubation. The C-terminal domain of PX thus appears to bind to the icosahedral assembly of fungal E2. To confirm the binding of the PX to the E2 interior, we collected single-particle cryo-EM data of this reconstituted tE2+PX recombinant subcomplex. The reconstruction shows an interior density which under icosahedral symmetry (i.e. when over-symmetrized) appears identical to that observed in the endogenous preparations (Fig. 3A, S6). The overall resolution is better, likely due to the absence of the PDC exterior. However, the improvement is localized to the E2 region, which we attribute to a decreased occupancy and order of the interior assembly compared to the endogenous preparation. The increased quality of the E2 region may also dominate alignment and increase implicit symmetrization, further de-emphasizing the interior component. These results indicate that the interior density observed in the native PDC reconstruction can be accounted for by the C-terminal domain of PX, namely the core-binding domain (CBD).

To identify the possible regions of the CBD responsible for interaction with the E2 bridge, we co-expressed His-tagged N-terminal PX truncations alongside an untagged E2 core-forming domain, inspired by a prior investigation^26^. Affinity purification on Ni-NTA resin indicated co-purification of E2 by PX-constructs omitting LD and PSBD, but including the full CBD (Fig. 3C, lanes 3-6). N-terminal truncations of PX beyond D280 appear to alter the interaction, despite equivalent expression of the PX-construct (Fig. 3C, lanes 7-12). This indicates that the CBD is required in its entirety to maintain its functional fold within the E2 core assembly, possibly affecting oligomerization and/or binding. Specifically, constructs PX301 and PX321 pulled down diminished levels of tE2. We attribute this to either decreased affinity for E2 or increased PX:E2 stoichiometry. Disruption of oligomerization may thus have caused an increase in bound PX by omission of the steric condition imposed by core-interior oligomerization. Truncation past G340 however abolished binding completely (cf. Fig. 3C, lane 2,10-12). This suggests that a critical binding motif is localized N-terminal to G340.

### 1.3 Domain architecture of the CBD

To further map the CBD, we identified 341 homologs of PX from various fungal species. Most share the expected domain topology LD-PSBD-CBD, while some Eurotiomycetes sequences lack the LD (Fig. S7). The sequence similarity in the CBD indicated a suitable clustering of Pezizomycotina (Pez) separate from Saccharomycotina (Sac), and a small set of dissimilar sequences. The latter set was omitted, and separate alignments were made for Pez (including *N. crassa*, 260 sequences) and Sac (including *S. cerevisiae*, 61 sequences). Alignment of Pez sequences indicated seven conserved prolines P1-P7 (Fig. 3D, S7), and a proline-rich region of lower conservation between P5 and P6. Here, a stretch of eight conserved residues form a DJJD*ϕ*LxG-motif (J=I/L, *ϕ*=hydrophobic, x=any) with the consensus sequence DIFDLLAG (Fig. 3D, motif M3). This motif was predicted as possibly helical, and could correspond to an acidic BH complementary to the basic/hydrophobic H2_E2_ pocket (Fig. 2). Notably, alignment of Sac sequences revealed the same motif. Alternatively, two similar conservation motifs are found in Pez, just before P3 and just after P4 (Fig. 3C and S7C, motifs M1-2), although their charge is less conserved. N-terminal truncation of the second of these motifs coincides with abolished binding (Fig. 3C). Definitive assignment of the BH will however require more detailed biochemical and/or structural investigation.

One long helix was predicted between P6 and P7 (Fig. S7C), likely comprising part of the OD observed by cryo-EM. Interestingly, beta strands were predicted and corroborated by a strong co-evolutionary signal, indicating a two-strand sheet (Fig. S7). While there was no obvious correspondence between the reconstructed OD and the predicted secondary structure elements, a co-evolutionary analysis of the Sac-alignment indicated a similar and partially coinciding three-strand sheet. Together with the conservation of the M3 motif (DJJD*ϕ*LxG) in sequences of otherwise low similarity, our observations indicate that i) the M3 motif is a likely candidate for the BH that is shared across ascomycota, and ii) the overall fold of the OD and/or entire CBD may differ in *N. crassa* vs. *S. cerevisiae*. It is also possible that multiple motifs from each PX monomer occupy multiple binding sites to limit binding stoichiometry, however we see no conclusive indication of this in our cryo-EM reconstructions.

### 1.4 *N. crassa* PX is essential for PDC function

To confirm the impact of PX on glucose metabolism, we assayed activity of PDC purified from wildtype and PX-knockout *N.crassa*. We first confirmed that wildtype PDC requires pyruvate, NAD and CoA for activity, as well as supplemented cofactors Thiamine pyrophosphate (TPP) and Mg^2+^. Enzymatic activity to 0.5 *μ*mol NADH min^*−*1^mg^*−*1^ was two orders of magnitude higher than reported for reactivated mammalian tissue extracts^27^, but an order of magnitude lower than what has been reported for PDC isolated from cauliflower^28^ or *Zymomonas mobilis*^29^. Conversely, PDC purified from the PX-knockout showed a complete loss of activity (Fig. S4). Mass spectrometry confirmed the absence of PX and E3, suggesting that the loss of PDC activity is due to disrupted E3 recruitment. In a differential growth assay, knockout growth on 1% Naacetate was similar to wildtype, but nearly ablated on 1% sucrose (Fig. S8). Inhibited growth on sucrose but not Na-acetate indicates a metabolic dysfunction attributable to impaired PDC function. This indicates that the PX-knockout relies on free mitochondrial E3, which drastically impairs pyruvate metabolism through the PDC *in vivo*. Wildtype and knockout strains showed comparable E1*α*-phosphorylation of S317, which has been shown to deactivate the human PDC^12,30^(Human-S293, or S264 in some references). This indicates that suppressed growth did not arise from differences in kinase/phosphatase recruitment.

### 1.5 Cryo-EM data is hidden by marginalized classification

The *N. crassa* PDC core is rigid and consequently its reconstruction is more reliable than the exterior, where the convolution of reconstruction is instead dominated by the distribution of the flexible component positions. Direct interpretation is therefore not possible in this part of the reconstruction. At most, elevated density proximal to the two- and five-fold symmetry axes of the dodechadral core (cf. Fig. 1) could indicate a predominant position for E1 and/or E3 since such elevated density is not observed along any three-fold axis, even under faithful reconstruction of tetrahedral symmetric classes. Interior to the core, PX oligomers have specific interactions that reduce flexibility and thus produce faithful reconstructions under appropriate symmetry, alignment and classification. However, the appropriate use of symmetry is ambiguous in terms of a faithful and productive classification. For instance, we observe a portion of the data to be clearly over-symmetrized during T-ASU classification, but to then reconstruct as pseudo-tetrahedral when the enforced symmetry was subsequently *lowered*. We attribute this to the increased ability to marginalize (spread) and subsequently de-emphasize conflicting data under lower symmetry. The classification outcome is thus clearly dependent on the extent to which the reconstruction procedure is able to marginalize data, a factor which is natural but, to the best of our knowledge not previously noted or quantified. By consequence any quantitative inference of classification demands consideration of the extent of marginalization by (but not limited to) symmetry. In the present example, it appears that some portion of particles must be hidden in order for a small portion of tetrahedral or near-tetrahedral particles to be distinguished from an over-symmetrized reconstruction. In a more general consideration, marginalization provides a reservoir for conflicting data in the form of an incoherent background, and its capacity depends on the extent of the space across which we permit marginalization.

To illustrate the capacity of marginalized reconstructions to hide data in this way, we recombined data pertaining to the distinct tetrahedral arrangements X and Y, in equal proportion, and ran a tetrahedral 3D-refinement with a mask made from the union of masks covering either tetrahedral arrangement. This reflects incompletely classified data with similar but irreconcilable sub-populations. We find that one of the arrangements overpowers the other, as the reconstruction is assuredly not the superposition of both arrangements (Fig. S9). This occurs despite the core dominating the alignment so that marginalization of the overpowered population is limited to 5 degenerate views within the T-ASU. Even so, one class apparently dominates completely. The superposition of the classes be recovered only when marginalization is entirely eliminated through reconstruction using alignments established for each class individually. This is thus a strong indication that marginalized reconstruction risks hiding large portions of data, and that it is consequently impossible to directly interpret any class population which has not been validated by other methods.

## 2 Discussion

The cryo-EM reconstruction of the *N. crassa* PDC core exhibits strict icosahedral symmetry, but with an additional protein PX that forms a tetrahedral assembly interior to the core under optimal conditions. Two distinct tetrahedral configurations could be identified and refined to 4Å resolution. Despite a completely different structural mechanism, we confirm that the fungal PX-component mirrors the mammalian E3BP as a selective recruiter of E3. Rather than substituting core E2 domains, PX oligomerizes interior to the PDC core (Fig. 4), occupying a significant portion of its volume. The oligomerization and volume occlusion by PX naturally suggest a mechanism for the variety of binding stoichiometries previously found for *S. cerevisiae*. We show conclusively that throughout ascomycota, PX binding stoichiometry is limited by the 30 available binding sites, located in the twofold symmetric interface connecting E2 core trimers. In *N. crassa*, the PX C-terminal domain oligomerizes into trimers. Oligomerization of PX limits binding stoichiometry to at most 24 PX monomers per E2 core as a result of PDC core geometry (cf. Fig. S1), and results in tetrahedral symmetry under favorable arrangement (Fig. S1, config 8S). The size of the PX oligomers also limits the available configurations through volume occlusion, so that the *N. crassa* E2 core can accommodate at most 12 interior PX monomers. Saturation of such optimal arrangement of PX oligomers again result tetrahedral symmetry of the PDC core interior. Volume occlusion does however degenerate the optimal arrangement into two distinctly different, tetrahedral arrangements (Fig. S1, config X4S and Y4S). We find *∼*60% of natively purified *N. crassa* PDC particles to be complete in this sense, but also observe directly that proportions of input data classified in conjunction with marginalizing reconstructions are not reliable indicators of the true distribution of states. Classes X and Y are however likely to dominate in the present sample, by evidence of the re-emergence of pseudo-tetrahedral reconstructions under lower symmetries and their correspondence to the geometrically optimal arrangements of interior density. Furthermore, threefold oligomers without occlusion should result in a tetrahedral arrangement that we see no evidence of. PX assembly and volume occlusion could also vary in related species, depending on the precise fold, oligomerization, or binding partners of interior PX components, such that other symmetries may be more prevalent in related fungal PDCs.

**Figure 4:**
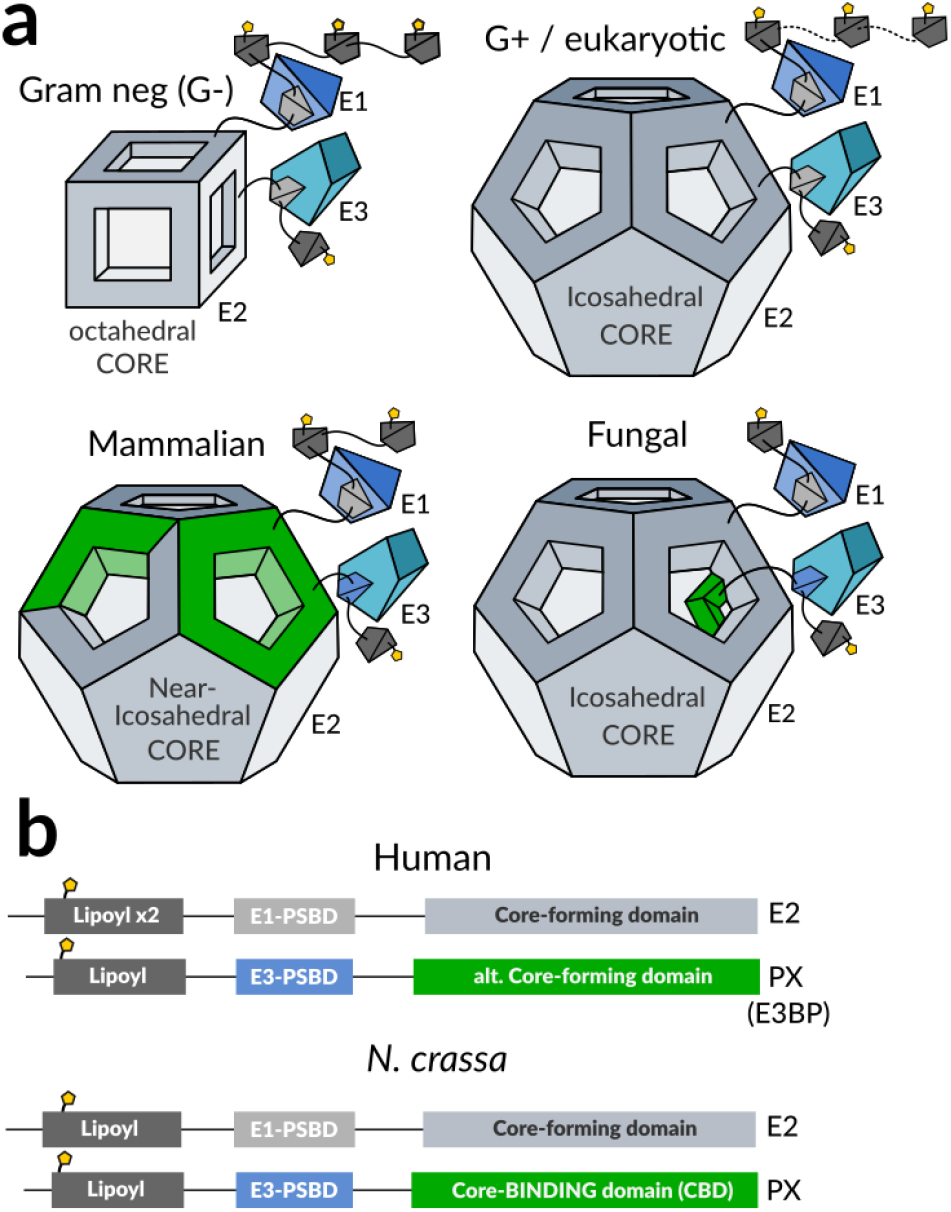
The fungal PDC is analogous in function to the mammalian PDC, but structurally distinct. **a**, Previously known PDC assemblies are octahedral and icosahedral core assemblies surrounded by flexibly tethered enzymes. The mammalian PDC has been shown to include an alternate core-forming protein which specifically recruits E3, but where an unknown mechanism and/or arrangement supplies a stoichiometric regulation to 40:20 or 48:12. The fungal PDC appears to also recruit E3 separately, but through a mechanism that maintains a homomeric core with strict icosahedral symmetry. The additional protein in the case of *N. crassa* forms a tetrahedral assembly interior to the core under optimal conditions, which results in a binding stoichiometry in direct correspondence to the icosahedral and tetrahedral symmetries, i.e. 60:12. The fungal PDC thus appears to have arrived at a similar stoichiometry and functional equivalence as the mammalian PDC, through an entirely distinct structural mechanism. **b**, The domain topology and function of mammalian and fungal PX are strikingly similar, despite their disparate structural rationales.

N-terminal truncation of PX to include only the CBD has shown a binding of 30 PX per core in *S. cerevisiae*^20,24^, consistent with monomeric PX binding. A reduction to 24 PX per core was observed if both the PSBD and CBD are left intact. Using our structural results we can now attribute the latter to binding of a trimeric PX oligomer without volume occlusion as depicted in Fig S1F. Our observations of altered binding affinity or stoichiometry in PX301 and PX321 (Fig 3C) and indications of a long-range beta-sheet interaction (Fig. S7) corroborate that the full CBD is required for oligomerization but not for binding. An earlier report showed that reconstituted PDC incorporating E1 and E3 bound only 12 PX per core^19^, consistent with volume-occluding trimers as in *N. crassa*. Volume occlusion by the PX oligomer may thus be further reliant on E1 and/or E3 binding, at least in *S. cerevisiae*. This has been rationalized by way of E3 docking into the pentagonal faces of the E2 scaffold^20^, a mechanism which seems unlikely given the PSBD of PX and the need for CoA to diffuse into the core interior. Moreover, we find no evidence of E3 docking directly to E2 in the native PDC from *N. crassa*. More likely, E1 and/or E3 might stabilize the PX trimer or contribute to its volume occlusion by binding to it. This might also explain why classification of recombinant E2-PX subcomplexes consistently fails to identify a tetrahedral core interior similar to the native *N. crassa* PDC in our hands, despite the obvious occupancy of PX in the core interior (Fig. 3A).

If PX can bind in a monomeric form when oligomerization is disrupted^21^, it remains unclear why oligomerization of the CBD with or without volume occlusion prevents binding of additional monomeric PX. We first consider that PX might be an obligate oligomer in its full-length form. This however seems unlikely, given the limited contacts made within the oligomer and the inability of PX oligomers to fit through the pentagonal openings of pre-formed core assemblies. It seems more likely that PX oligomerizes upon binding to the core. We also observe fast incorporation of PX30 into the PDC core during subcomplex reconstitution, further supporting a model in which PX oligomerization and binding are mutually enhancing. So what prevents monomeric PX from binding in excess of oligomerized PX as indicated by previous investigations? The identification of multiple similar conserved motifs in Pezizomycotina consistent with the core binding pocket (Fig. S7, motifs M1-3) could suggest that each PX monomer persistently occupies multiple binding sites, however this does not explain previous observations in *S. cerevisiae* since only the M3 motif (DJJD*ϕ*LxG) is found in Saccharomycotina. Instead, we suggest that oligomerization may be required for retained ability to bind when the dynamics of the extended N-terminal domain(s) are intact. This is appealing as a mechanism of PDC regulation through fold stability and possible ligand and/or substrate interactions, and is consistent with all previous observations.

Finally, we note that *∼*40% of the endogenous PDC particles did not classify as containing either of the two tetrahedral arrangements of core-interior PX. This is most likely due to an unsaturated interior, but we cannot exclude the possibility of asymmetric interior due to sub-optimal arrangements in these particles. Each PX monomer binds to an interface formed by multiple core trimers, so that PX can only bind partially or completely assembled cores. This agrees with the notion of monomeric PX which oligomerizes upon PX binding to access the core interior, and is supported by our observations of fast incorporation of PX into assembled E2 cores following separate expression. Interestingly, in a naive model of assembly in which trimeric PX oligomers attach sequentially to a pre-existing core, only 15% would be expected to be in each tetrahedral arrangement - half of what is indicated by our classified proportions of cryo-EM particles. This discrepancy could possibly be attributed to uncertainty of classification due to possible data hiding. However, if the classification is accurate, it would suggest that PX co-assembles with the PDC core by binding to core fragments, which leads to an enrichment of optimal final arrangements and thus higher efficiency.

## 3 Methods

### 3.1 N. crassa cultivation and mitochondrial purification

Frozen *N. crassa* (WT: STA74 / ATCC 14692, KO: FGSC 15821) mycelium was thawed and inoculated on 50 ml 2% agar minimal (Vogel’s) medium supplemented with 1.5% sucrose and 1.5% Na-Acetate in 200 ml flasks. After incubation for 72h at 30^*◦*^C in darkness followed by 72h at room temperature in daylight, or after conidia appeared, conidia were harvested and filtered using sterile water. 500 ml Vogel’s medium supplemented as above were inoculated with conidial suspension, and grown in an flask (2L) for 24 h at 25^*◦*^C under illumination and shaking at 140 rpm. 25 ml of the resultant growth were then used to inoculate each of 20 E-flasks (2L) containing 500 ml Vogel’s medium, and then grown for 16 h at 25^*◦*^C under illumination and 140 rpm shaking. Subsequent steps were conducted on ice, or at 4^*◦*^C. 50 g of Mycelium was collected by filtration through double-layer muslin, and ground using a chilled mortar, adding 100 g SiO_2_ sand and 50 ml Sucrose-EDTA-MOPS (SEM) buffer containing 0.5% phenylmethylsulfonyl fluoride (PMSF). Sand was pelleted by centrifugation at 2000 rcf for 10 min, collecting supernatant. The pelleted sand and cellular material mixture was ground twice more and pelleted, each time adding fresh buffer and pooling supernatant. Differential centrifugation then followed; 17,500 rcf for 20 min, collecting pellet. The pellet was resuspended in 30 ml SEM and clarified at 2000 rcf for 10 min. Crude mitochondira were then pelleted at 17,500 rcf for 20 min, and resuspended 10 ml SEM. The re-suspended crude mitochondria were loaded on a 15-23-32-60% (4.5-4.5-12-4.5 ml) sucrose gradient and centrifuged in a Beckmann SW28 swing-out rotor at 100,000 rcf for 1h. The mitochondrial band was collected from the 32-60% interface and frozen at −80^*◦*^C.

### 3.2 Endogenous PDC purification

Mitochondria were thawed on ice and lysed using DDM to a final concentration of 2%, incubated under mild rocking for 15 min. Non-solubilized matter was pelleted and removed at 30,000 rcf for 20 min. The clarified solution was loaded onto one or more 1M sucrose cushions (7 ml) in Ti70 tubes and the sample pelleted in a fixed-angle rotor using 230,000 rcf for 4h. Pelleted material was gently washed and resuspended in 0.4 ml of 50 mM HEPES-KOH (pH 7.5), 20 mM KCl, 1 mM DTT, then clarified at 13,000 rpm for 10 min. The clarified solution was loaded onto a continuous 20-40% sucrose gradient in SW40 tubes and separated in a swing-out rotor at 70,000 rcf over 16h. PDC particles formed a visible band that started rather abruptly at 33% sucrose, extending with a smooth decrease towards higher sucrose concentrations. Fractions containing PDC were collected from the gradient using a long-needle syringe. This produced pure and intact PDC particles within 20 hours of mitochondrial lysis. Buffer exchange and purification was conducted in two steps. First, material was pelleted in TLA120.2 tubes at 100,000 rcf for 2h, resuspended in 0.25 ml fresh buffer, and twice clarified at 13,000 rpm. Finally, was fractionated using size-exclusion chromatography (SEC) (GE Superose 6-increase 3.2/100).

### 3.3 Recombinant expression and purification

The following genes were purchased from Life Technologies Europe for bacterial expression of *N. crassa* genes: E2 (29-458 uniprot:P20285), tE2 (225-458 uniprot:P20285), PX (30-426 uniprot:Q7RWS2). The pRSET vectors were used for single expression and provided a His6-tag that is cleavable using either thrombin (E2,tE2) or TEV (PX). petDuet dual-expression vectors were constructed using these as template, in all cases positioning the PX gene in the His-tagged MCS1. All vectors were amplified using *E. coli* DH5-*α*, and all expression utilized Rosetta2 (DE3). For expression, cells were grown in terrific broth (TB) at 37^*◦*^C and 180 rpm until OD reached approximately 0.5. Expression was induced by addition of IPTG to 1mM and allowed to proceed for 3h before harvest. Expression was consistently found to be worse if conducted for 16h at 18^*◦*^C.

Cells were pelleted and resuspended in 50mM imidazole buffer, then lysed either through sonication (small-scale, e.g. truncation affinity assay) or high-pressure homogenization (large scale, cryo-EM preparations). Intact cells and debris were pelleted, and the supernatant collected. Ni-NTA agarose slurry was added and incubation under agitation proceeded for at least 30 min. Isolation and washing of Ni-NTA was performed either through repeated pelleting (small scale), or a gravity flow column. Depending on the downstream purpose, protein was either cleaved using TEV or eluted using imidazole. E2 was found to be well-expressed in all examined forms. Expression of PX was found to be much lower, but greatly improved in the presence of co-expressed E2 or tE2.

### 3.4 Resolving the Knockout homokaryon

A NCU00050-knockout strain was purchased from the FGSC (15821), available as a heterokaryon with a wildtype helper genome to maintain viability on standard media. We resolved the KO homokaryon by generating monokaryotic spores on crossing media^31^ with 2% sucrose and 1.5% agar. Microconidia were purified by filtering through double-layer 5V Whatman filters. The presence of microconidia and the absence of macroconidia was confirmed by microscopy. Spores were diluted appropriately and plated on fresh crossing media containing 40 mM acetate, 1% sorbose (to promote colonial growth), 1.5% agar, and 300 *μ*M hygromycin (selecting for the knockout genome). The use of acetate in place of sucrose enabled a KO homokaryon to survive. 10 colonies were picked after a few days and replated on 5 ml fresh Vogel media plates, using 40mM acetate, 1.5% agar, and 300 *μ*M hygromycin, and grown until spores were produced. Spores were again purified and next subjected to a differential growth assay in Vogel’s media, where a homokaryon knockout was expected to grow more favorably in 40 mM acetate compared to 40mM sucrose. Cultures for two out of the 10 colonies followed this expectation. The above procedure was repeated for one of these 2 cultures to ensure complete purity, supplementing the crossing media with 40 mM Na-acetate. The subsequent round of the differential growth assay showed 10 out of 10 colonies growing better on Na-acetate than sucrose. One of these colonies was used for subsequent cultivation and purification.

### 3.5 Activity assay

The activity of intact PDC assemblies was assayed by time-dependent spectroscopic monitoring of NADH production at 340 nm and room temperature. All samples for activity measurements were purified by SEC. The reaction was conducted in a UV-transparent (Sigma-Z628026) cuvette, with reactants as described in table 1.

**Table 1:** Reactants in activity assay.

| | Stock [mM] | Use [μl] | Final [mM] |
| --- | --- | --- | --- |
| HEPES pH7.5 | 20 | 800 | 17.0 |
| NAD | 50 | 50 | 2.5 |
| Na-Pyruvate | 100 | 20 | 2.0 |
| Na-CoA | 10 | 20 | 0.2 |
| TPP | 10 | 20 | 0.2 |
| MgCl | 50 | 20 | 1.0 |
| PDC | 2.2 mg/ml | 10 | 23 μg/ml |
| Total |  | 940 |  |

All components apart from pyruvate and purified PDC were added and mixed, and a baseline was collected. Purified PDC was added at t=−1 min. At t=0 min the reaction was initiated by addition of pyruvate. The activity decreased with time and was therefore resolved in time covering a 30 s sliding window, and linearly extrapolated to t=0. At no stage of growth, purification or assay was phosphatase or dichloroacetate introduced to promote dephosphorylated E1.

### 3.6 Mass spectrometry

Peak fractions following SEC were immediately frozen and used for mass spectrometry. Protein identification and quantification were carried out by the Proteomics Biomedicum core facility, Karolinska Institutet. Analysis was performed using Scaffold and Scaffold PTM (post-translational modification).

### 3.7 Cryo-EM grid preparation and data collection

Grids for cryo-EM were prepared by glow-discharge in a Pelco easiGlow. 3 *μ*l of sample was then applied to the grid and vitrified in a FEI Vitrobot mark IV, following 30 s wait, 2 s blot, and 2 s additional wait before plunging. Grid screening and optimisation, as well as data collection was conducted at the Swedish National Cryo-EM Facility at SciLifeLab, Stockholm University and Umeå University. The parameters of grid preparation and data collection are summarized in table 2.

**Table 2:** Datasets collected. Calibration of magnification was performed for reconstructions with associated atomic models. For tE2+PX30 calibration was performed by PHENIX-refinement against multiple rescaled maps. The best agreement in terms of overall refinement statistics was chosen. For native, the map was rescaled to correlate maximally with the calibrated tE2+PX30 map.

| Dataset | Native | E2 | tE2+PX30 | tE2/PX261 |
| --- | --- | --- | --- | --- |
| Grid |  | Quantifoil R1.2/1.3 | Quantifoil R1.2/1.3 | Quantifoil R1.2/1.3 |
| Carbon support | Yes | No | No | No |
| Microscope | FEI Talos | FEI Talos | FEI Krios | FEI Krios |
| Detector | Falcon 2 | FEI Falcon 3 | Gatan K2 GIF | Gatan K2 GIF |
| Voltage | 200 kV | 200 kV | 300 kV | 300 kV |
| Magnification |  |  |  |  |
| Nominal | 114k (1.23Å/pix) | ( 1.23Å/pix ) | 165k ( 0.83Å/pix ) | 165k ( 0.86Å/pix ) |
| Calibrated | 112k (1.25Å/pix) | - | 163k ( 0.85Å/pix ) | - |
| Mode | Counting | Linear | Counting | Counting |
| Gain normalized | Yes | Yes | Yes | Yes |
| Micrographs | 4867 | 344 | 5675 | 4063 |
| Total dose | 35.0 e/Å<sup>2</sup> | 33.5 e/Å<sup>2</sup>, 4s | 27.3 e/Å<sup>2</sup>, 4s | 31.4 e/Å<sup>2</sup>, 4s |
| Fractions | 19 | 39 | 20 | 32 |

### 3.8 Cryo-EM processing

Preprocessing was conducted using motioncor2^32^ and Gctf^33^. All reconstruction, alignment, classification and postprocessing was conducted using RELION^34^. The parameters of data processing and 3D reconstruction are summarized in table 3 and described in brief in the following sections.

**Table 3:**
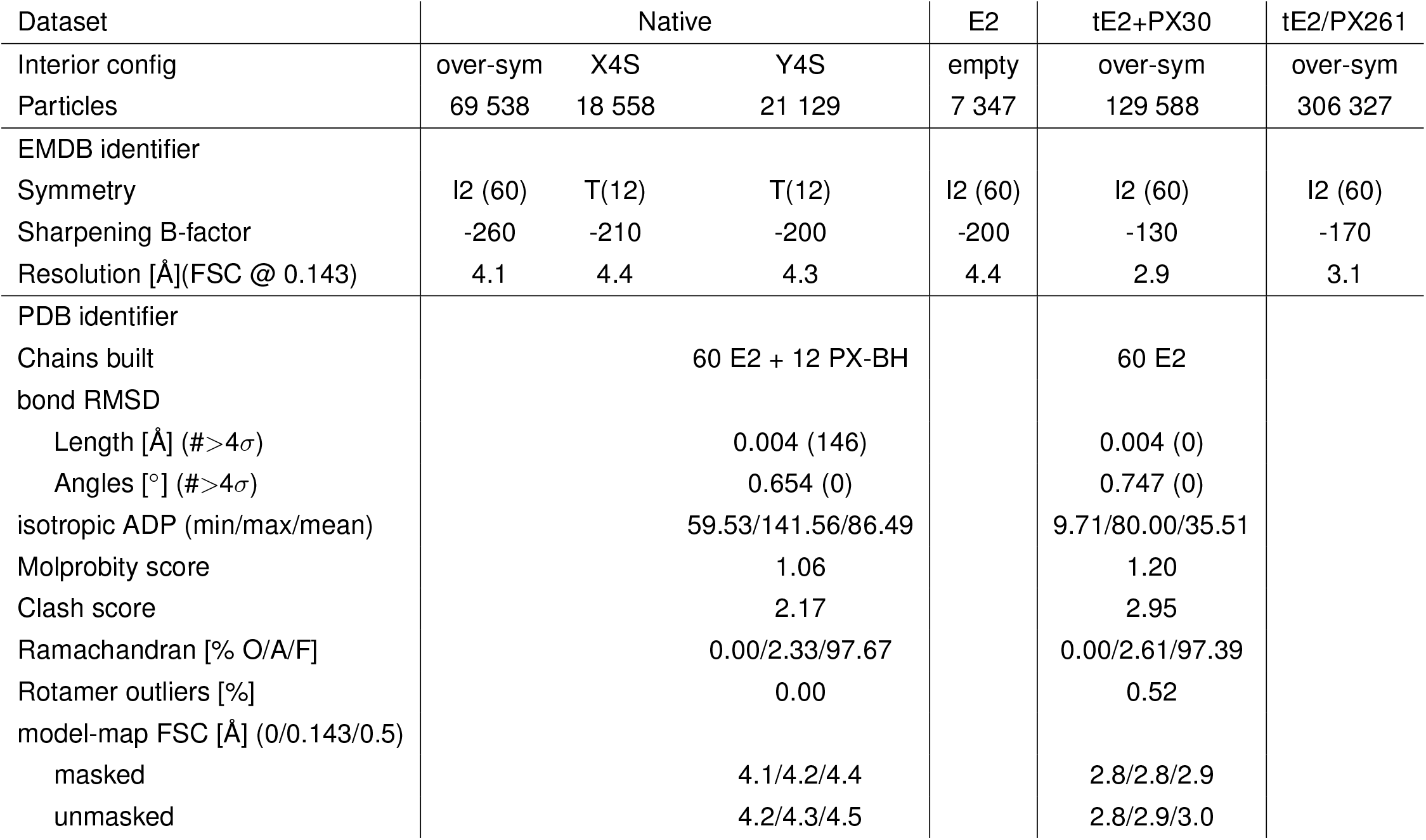
Reconstructions. tE2+PX30 refers to reconstitution of separately expressed components, whereas tE2/PX261 was a subcomplex of co-expressed components. The E2 core was built ab-initio into the PX30 reconstituted subcomplex reconstruction. The model for the native Y4S model was based on the same E2-core model, placing further small helical poly-Ala section into the density corresponding to the PX-BH. The records of these residues were changed to UNK **after** refinement, in line with recommendations.

#### 3.8.1 Native preparation

139 PDC particles were picked manually from nine micrographs and aligned by 2D-classification. The resultant class average was used to pick 350k candidate particles from 4867 micrographs. A subset of 139k particles was selected based on two sequential rounds of 2D-classification. An icosahedral initial reconstruction was generated in RELION. An icosahedral refinement was next performed under icosahedral symmetry to center particles. The magnification was adjusted, determined by rescaling the map to correlate maximally with the reconstituted E2-PX30 subcomplex which was used to build the atomic model.

Following re-extraction, a 3D-classification of 10 classes under icosahedral symmetry was used to further identify 70k particles that appeared to be of higher quality, based on visual inspection. This subset was refined to 4.1 Å (map N1), and used for all subsequent analysis. Classification identified two tetrahedral and one over-symmetrized class, used as seeds for a multi-reference classification (described below). Subsets of 30% each of the tetrahedral classes were found and individually refined to 4.2 Å and 4.3 Å respectively. As post-processing appeared to over-sharpen the lower-resolution interior, local-resolution filtering was conducted in RELION (maps N2X/Y).

#### 3.8.2 tE2+PX30 reconstituted subcomplex

225k particles were picked automatically from 5675 micrographs using WARP^35^. A consensus refinement was used to center and generally align particles. These (3D-)alignments where used to perform 2D-classifications without further orientation alignment, from which 213k particles were selected. These particles produced a 3.5 Å reconstruction under icosahedral symmetry. Following CTF-refinement and beam-tilt correction, the icosahedral reconstruction reached 2.9Å (map R1). The final magnification was adjusted by rescaling the map to optimize covalent geometry metrics during refinement in PHENIX. A tetrahedral 3D-classification using five classes identified a subset of 130k particles that were of higher comparative quality, as assessed by a 2D-classification of each subset, as well as a higher overall refined resolution (3.4 Å using icosahedral symmetry). Multi-reference classification identified 25% and 29% of these as tetrahedral PX-packing interior class X and Y respectively, which were refined to 3.2 Å and 3.6 Å under tetrahedral symmetry (map R2X/Y).

#### 3.8.3 E2 core

134 particles were picked manually from nine micrographs. These were averaged by 2D-classification and used as references for automated picking in RELION. Almost 25,000 particles were picked from 344 micrographs, followed by cleaning through multiple rounds of 2D-classification. 3D-classification under icosahedral identified further data which did not appear to represent E2-core particles. 7347 particles belonged in classes representing intact PDC core particles. These were refined to 5.4 Å, which after refinement of CTF and beam-tilt improved to 4.3 Å (map L1).

### 3.9 Model building

A molecular model of *N. crassa* E2 was built into EMDB-xxxx (map R1), covering residues 227-458. This model is deposited as PDB-xxxx. Additionally, the backbone trace of the binding helix of PX was built into EMDB-xxxx (map N1Y). This model is deposited as PDB-xxxx. Model validation metrics are shown in table 3. Molecular models were constructed using Coot^36^, and refined using PHENIX^37^. Images were produced using UCSF Chimera^38^ and chimeraX^39^.

### 3.10 Data availability

All reconstructions presented have been deposited in the Electron Microscopy Data Bank, and atomic models have been deposited in the Protein Data Bank. Accession codes are given in table 3.

### 3.11 Bioinformatics

Sequences annotated as ‘E3BD-containing’ were extracted from PFAM-32.0^40^ and subjected to a preliminary multiple-sequence alignment (MSA) in Jalview^41^. E2-like proteins were identified by presence of a conserved catalytic DHR-motif and/or preservation of secondary structure consistent with the known fold of E2, and omitted from further analysis. Alignment of this set of sequences formed the ascomycota PX MSA. The CBD was identified as everything following the flexible linker C-terminal to the E3BD-homologous domain, and isolated based on the PX MSA. Following a nearest-neighbor tree-clustering under BLOSUM62 substitution score, Pezizomycotina (Pez) and Saccharomycotina (Sac) were separated and re-aligned individually, forming the Pez-CBD and Sac-CBD MSA, respectively. Co-evolutionary correlation was conducted through the raptorX^42^ MSA-interface using the established CBD MSAs.

## 4 Acknowledgments

The presented research would not be possible without an initiative by Alexey Amunts, who we also thank for a lot of useful advice, suggestions and encouragement throughout the project. The cryo-EM data were collected at the Swedish national cryo-EM facility, staffed by M. Carroni, J. M. de la Rosa Trevin, J. Conrad, and S. Fleischmann. We also thank the members of Amunts and Lindahl lab for active discussions throughout the project.

## 5 Funding information

This work was funded by the Swedish Foundation for Strategic Research (FFL15:0325), the Ragnar Söderberg Foundation (M44/16), the Swedish Research Council (2015-04107 and 2017-04641), Cancerfonden (2017/1041), EU grant ERC-2018-StG-805230 and BioExcel-823830, the Knut and Alice Wallenberg Foundation (2018.0080), and the Lennander Foundation. It was also supported by the Knut and Alice Wallenberg Foundation, Family Erling Persson Foundation, and Kempe Foundations through the Swedish National Cryo-EM Facility.

## 6 Author contributions

BOF designed experiments, collected and processed data, and wrote the article. SA designed experiments and collected data. RJH designed experiments and wrote the article. NM contributed essential data and experimental procedures. AA suggested and supervised the project, and wrote the article. EL supervised data processing and wrote the article.

## 7 Competing interests

The authors declare no competing interests.

---

**Figure S1:**
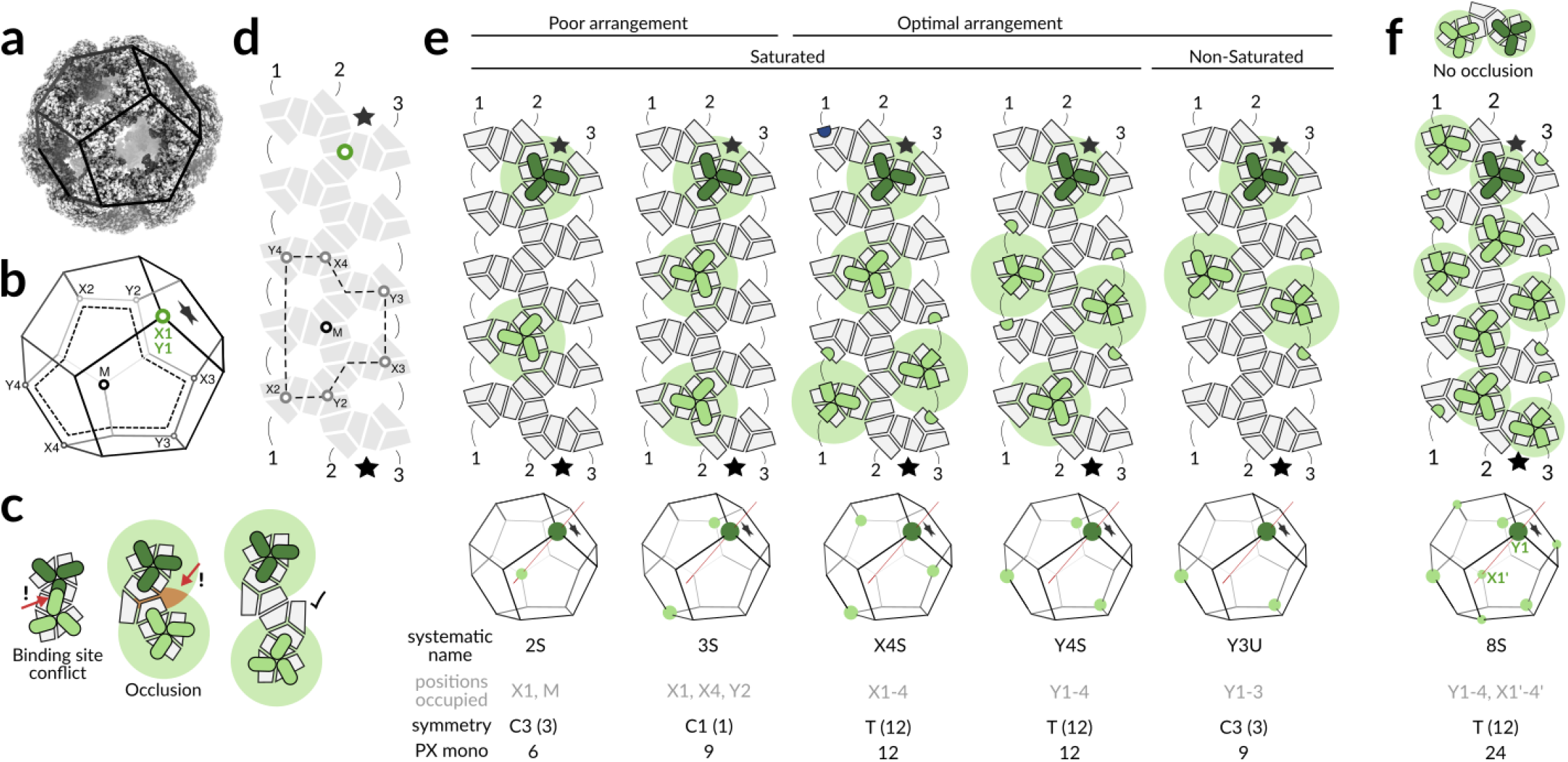
Configurations of the fungal PDC interior. **a**, The PDC core is a dodecahedron, where the vertices are E2 trimers, and edges are points of contact between adjacent such trimers. **b**, A sketch illustrates important vertices for the subsequent arrangement of PX interior to the PDC core. Each PX trimer occupies one E2 trimer in the sense that their threefold symmetry axes coincide and that the PX trimer binds all edges connecting the E2 trimer vertex. **c**, This means that adjacent vertices cannot be PX-occupied, since the edge is already occupied. The volume occlusion by PX also imposes a steric condition. This is illustrated schematically as a circular background centered on each PX trimer. It is evident from examination of 3D reconstructions that PX occupancy at vertices separated by exactly one intermediate vertex results in clashes of the PX oligomer-occluded volume. Rather, each PX-occupied vertex must be separated from any other by at least two edges. **d**, We ‘unfold’ the assembly of E2-trimers (which forms the PDC core) to easily illustrate **e**, the possible configurations of PX in the E2 core interior. The points corresponding to those in panel **b** are indicated, and one face is marked with a star for clarity. Numbered lines indicate edges that are broken by unfolding the dodecahedron into this 2D-schematic. Without loss of generality, we can choose one E2-trimer to always be PX-associated. All further consideration regards which additional E2-trimers may be occupied. There is exactly one configuration wherein just two PX trimers are placed such that no further PX trimer can be placed, hence saturated (2S, S=Saturated). We are able to construct at least one 3S configuration, and there are exactly two 4S configurations (X4S and Y4S). The latter two correspond to classes X and Y found in the native PDC data. We also show one example of an unsaturated configuration (Y3U, U=unsaturated), corresponding to a likely configuration in the non-tetrahedral class N. The resultant symmetry is also listed for each displayed configuration along with the order of the symmetry in parenthesis, as well as the number of PX monomers per core. **f**, If the oligomerization of PX interior to the core does not result in a volume occlusion as shown in panel **c**, the optimal arrangement is still tetrahedral, but with a doubled stoichiometry. This 8S configuration is equivalent to a superpositon of configurations Y4S and a rotated X4S.

**Figure S2:**
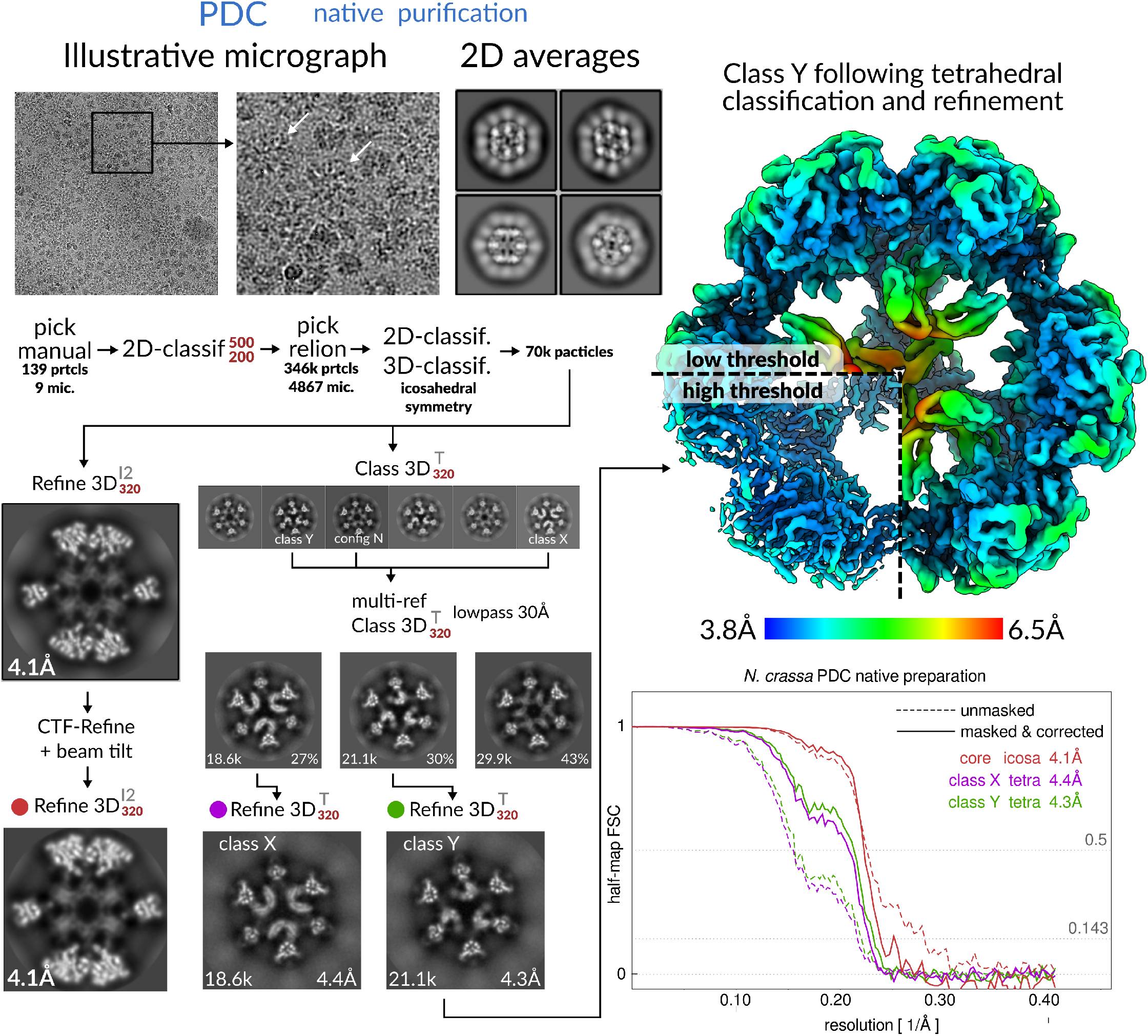
Processing of the native PDC. Multi-reference classification was conducted by providing three input references, corresponding to the three indicated class reconstructions X, Y and N. The full dataset (70k particles) was thus re-classified to assess the distribution among classes. Corrected FSC refers to the adjustment of the half-map according to over-fitting identified by phase-randomization beyond FSC=0.8.

**Figure S3:**
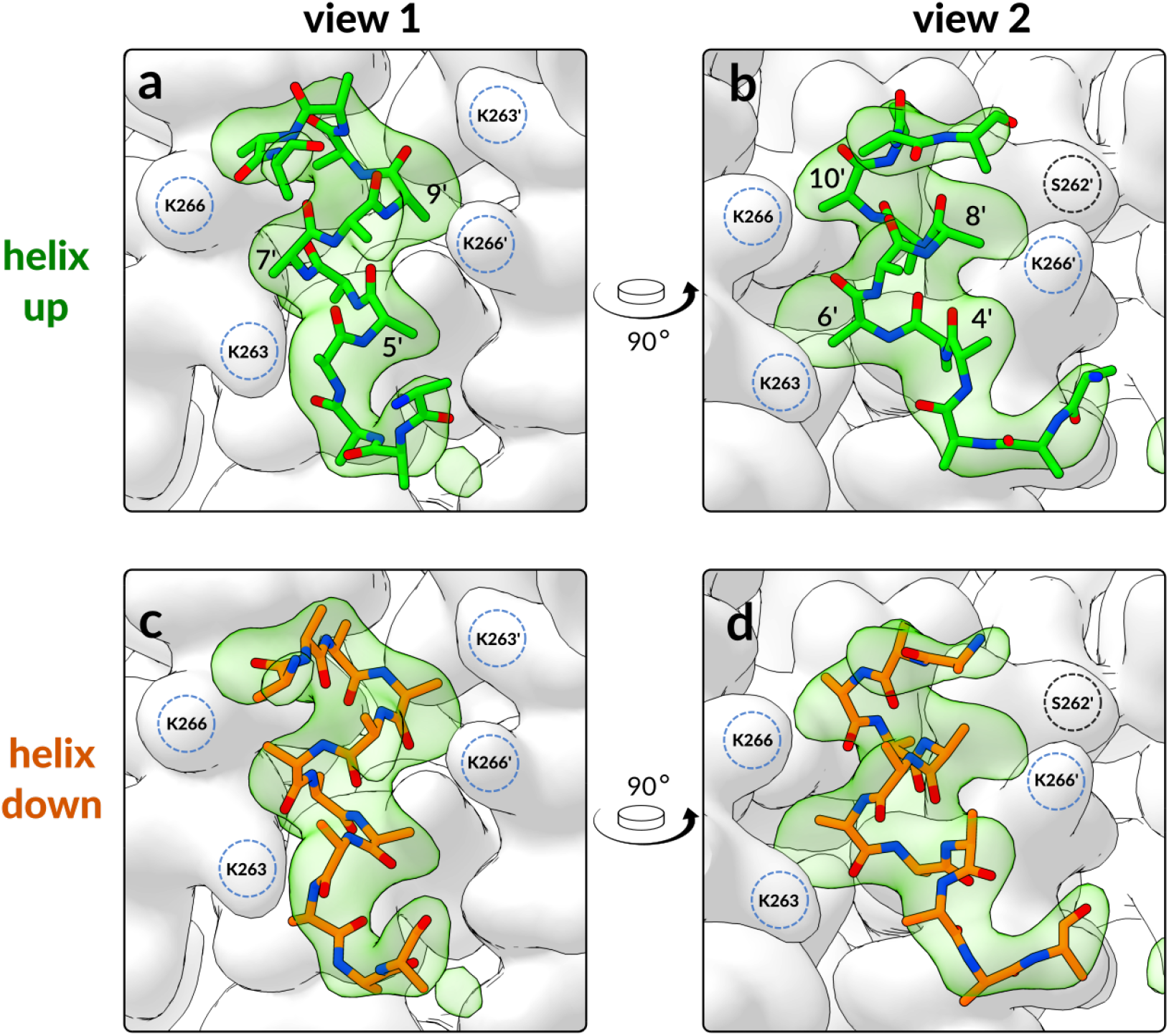
PX binding helix. The tetrahedral symmetry imposed to reconstruct the tetrahedral classes X and Y results in a break of the strict twofold symmetry of the E2 binding pocket where the PX binding helix (BH) binds, but the pocket itself which is composed of helix-2 (H2) of E2 still appears completely symmetrical, with no detectable induced fit. Even so, the BH appears to bind with a bias for one orientation, apparently enforced by the handedness of the oligomer. Its orientation is somewhat ambiguous, and so both possible orientations are built and considered. The limited resolution prevents modeling of side-chains, which in turn prevents a model-to-map cross-correlation from discriminating which orientation is more likely. However, based on the direction of weaker and residual side-chain densities, we nonetheless find one direction (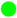 panels **a+b**), is more likely than the other (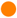 panels **c+d**). The density is identical in all panels. This interpretation leads to a stronger density connecting the BH N-terminus to the PX-oligomer, indicating that the BH is C-terminal to at least part of the oligomer domain.

**Figure S4:**
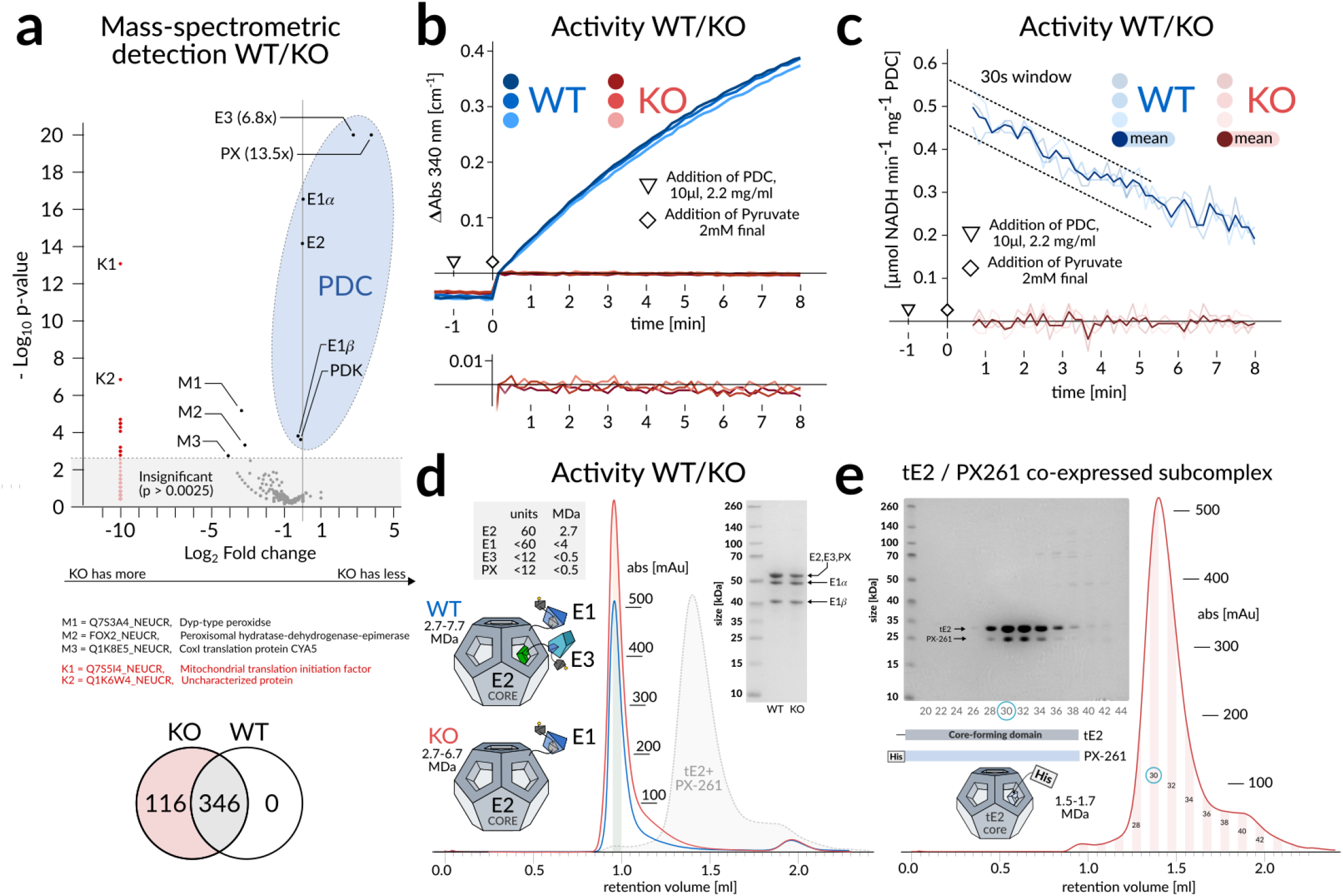
Size exclusion, activity and mass spectrometry. **a**, Mass spectrometry (MS) confirms that the knockout causes concomitant loss of E3, with no other changes attributable to the PDC-interaction. Some proteins appear to be up-regulated in response to PX-knockout. We cannot rationalize why these would be co-purified with the PDC through SEC, other than their general abundance and the sensitivity of the MS analysis. **b**, Time-resolved kinetics of the purified PDC confirms the detrimental effect of PX knockout on PDC function. Purified complex and pyruvate were added at *t*=−1 min and *t*=0 respectively, to a cuvette containing all other necessary reactants (see methods, table 1). **c**, Same as in panel **b**, but with the rate of change averaged over 30s windows following initiation of the reaction. PDC function appears completely eliminated in the KO, attributable (but not necessarily limited) to loss of E3 as shown in panel **a**. The activity appears to decrease rapidly, so the WT activity is extrapolated to *∼*0.5*μ*mol NADH min^*−*1^mg^*−*1^ PDC at *t*=0. **d**, Size exclusion chromatography (SEC) of wildtype and knockout PDC indicate similar native size, which is expected given the small relative contribution of PX and E3. An SDS-gel of the peak fraction (shaded area) displays a weakened band where E2, PX and E3 overlap. **e**, A co-expressed subcomplex shows that affinity-purified PX261 retains a high affinity to the PDC core, with no detectable unbound PX following SEC. This may indicate a stoichiometric deficit of PX261 and subsequently unsaturated core complexes, which could rationalize why tetrahedral classes are more elusive in this cryo-EM dataset, despite the obvious occupancy of the core interior. This is corroborated by the observation that expression of PX261 is greatly hampered in the absence of co-expressed E2 or tE2, indicating that it is sensitive to degradation or proteolysis.

**Figure S5:**
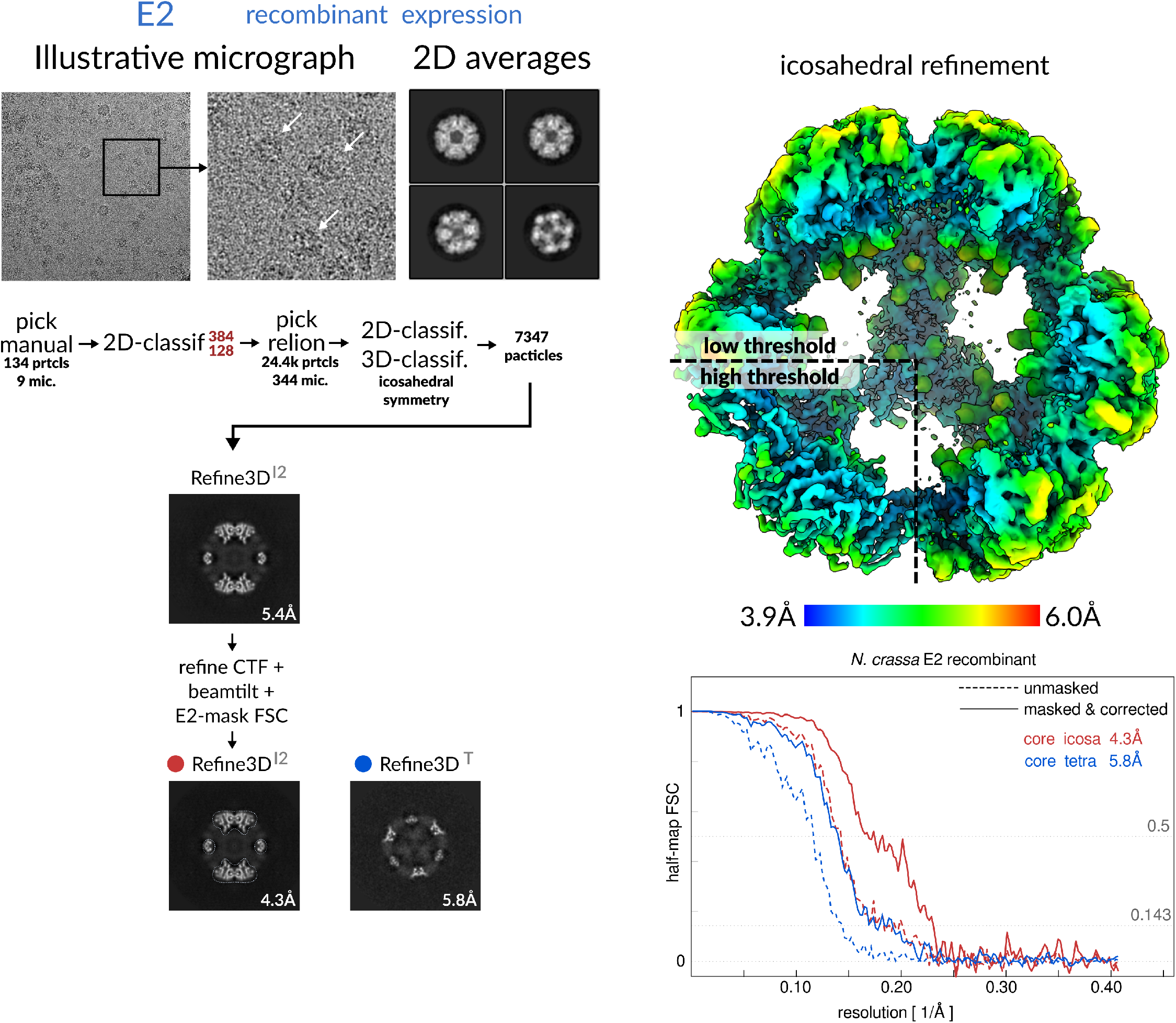
Processing of the E2 core assembly. Corrected FSC refers to the adjustment of the half-map according to over-fitting identified by phase-randomization beyond FSC=0.8

**Figure S6:**
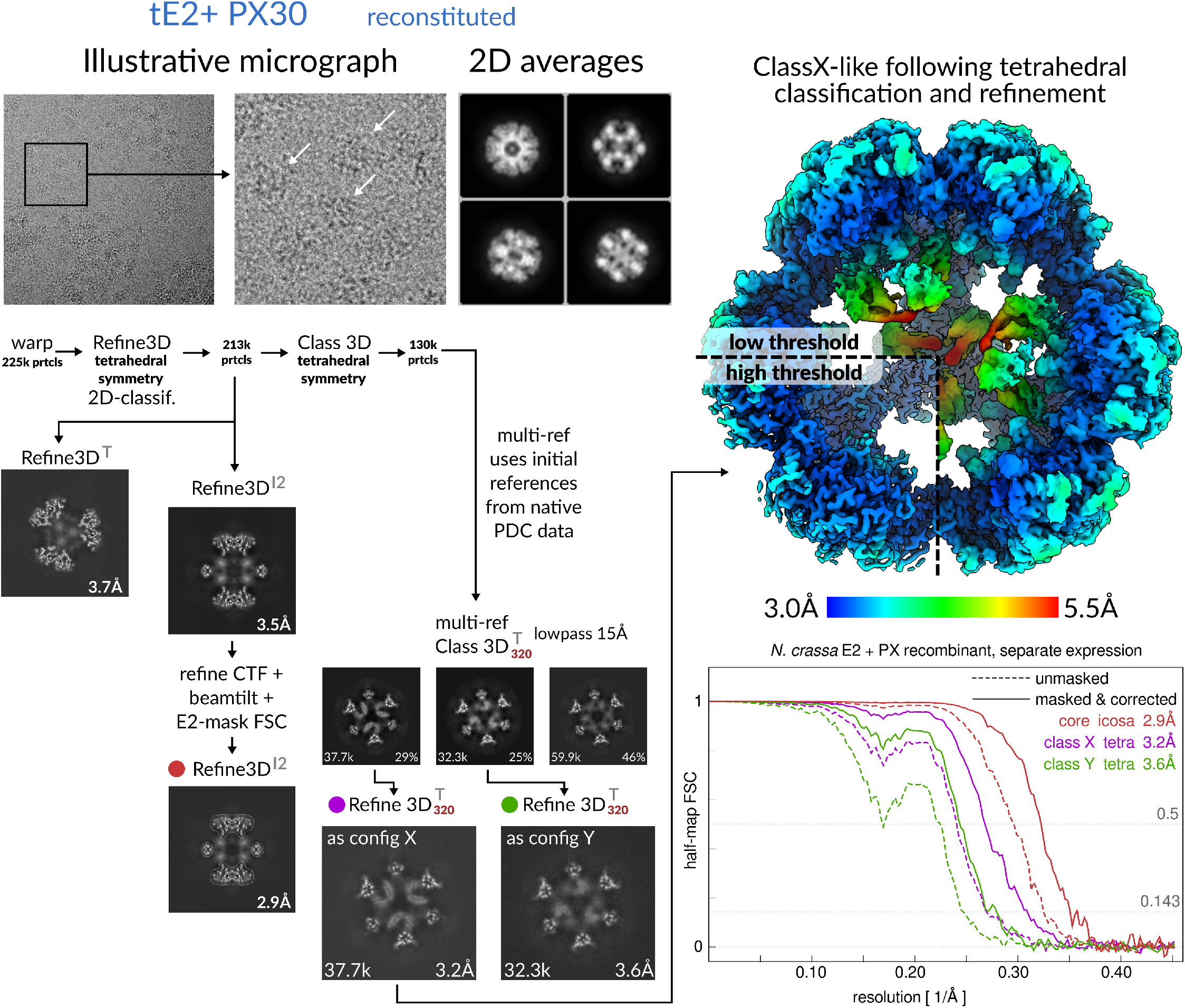
Processing of E2+PX subcomplex. Multi-reference classification was conducted by providing three input references, corresponding to the three indicated class reconstructions X, Y and N *from the native PDC classification*. This introduces a measure of reference-bias. The full dataset (70k particles) was thus re-classified to assess the distribution among classes, but no inference was made from this distribution. It is shown here for completeness and to indicate the correspondence of the interior density with PX trimers. It should not be taken to indicate correspondence of the interior arrangement with configurations X or Y as a whole. Corrected FSC refers to the adjustment of the half-map according to over-fitting identified by phase-randomization beyond FSC=0.8

**Figure S7:**
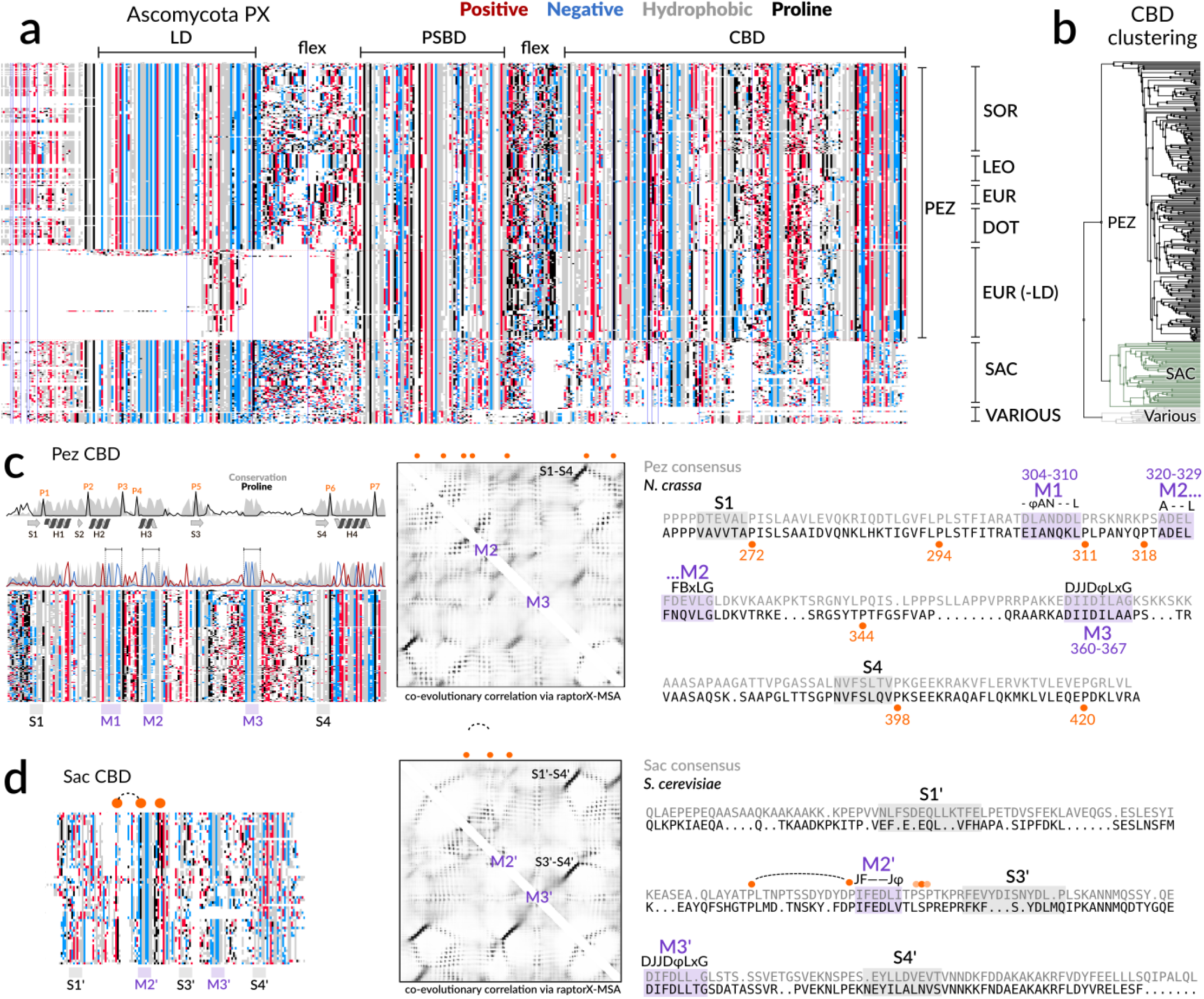
Multiple sequence alignemnts of fungal PX. **a**, The overall multiple sequence alignment (MSA) of fungal (ascomycota) PX, as found by fetching all PFAM sequences annotated as E3BD-containing (the current annotation for the PSBD), filtered to exclude any E2-like core-forming protein. Three-letter labels (SOR/LEO/EUR/DOT) refer to classes within the subdivision Pezizomycotina (PEZ). Notably, the lipoyl domain (LD) is conserved but missing in some EUR sequences. The LD is not well-aligned in the aberrant sequences (labeled ‘various’), casting doubt with regards to their purpose and/or annotation. **b**, Clustering of the isolated CBD shows PEZ and Saccharomycotina (SAC) as fundamental clusters. **c**, Isolating and re-aligning the Pez CBD (incl. *N. crassa*) shows strong conservation, especially of seven prolines that appear to delimit the overall domain topology. A co-evolutionary correlation (raptorX) using this MSA finds support for at least one long-range anti-parallel beta-sheet, supported by secondary structure prediction. There are also three similar and conserved negative motifs, M1-3. The consensus motifs are shown (J=I/L, B=D/N, Z=Q/E, *−*=D/E, *ϕ*=hydrophobic, x=Any). It is also notable that the region spanning P5 and P6 has low conservation but high charge and proline content. This region also contains M3. The PEZ consensus sequence is aligned to *N. crassa*, mapping P1-7 and the conservation motifs. Notably for *N. crassa*, M1 is deficient in the predicted negative charge and the P5-P6 region has surprisingly low proline-content. **d**, the SAC CBD shows lower conservation of both proline and overall residues, with the notable exception of the DJJD*ϕ*LxG-motif, which is shared with PEZ. A similar motif (JF*−−*J*ϕ*) is also found, which does not have an obvious correspondence to either of the other motifs found in PEZ, but which has an overall correspondence with M2.

**Figure S8:**
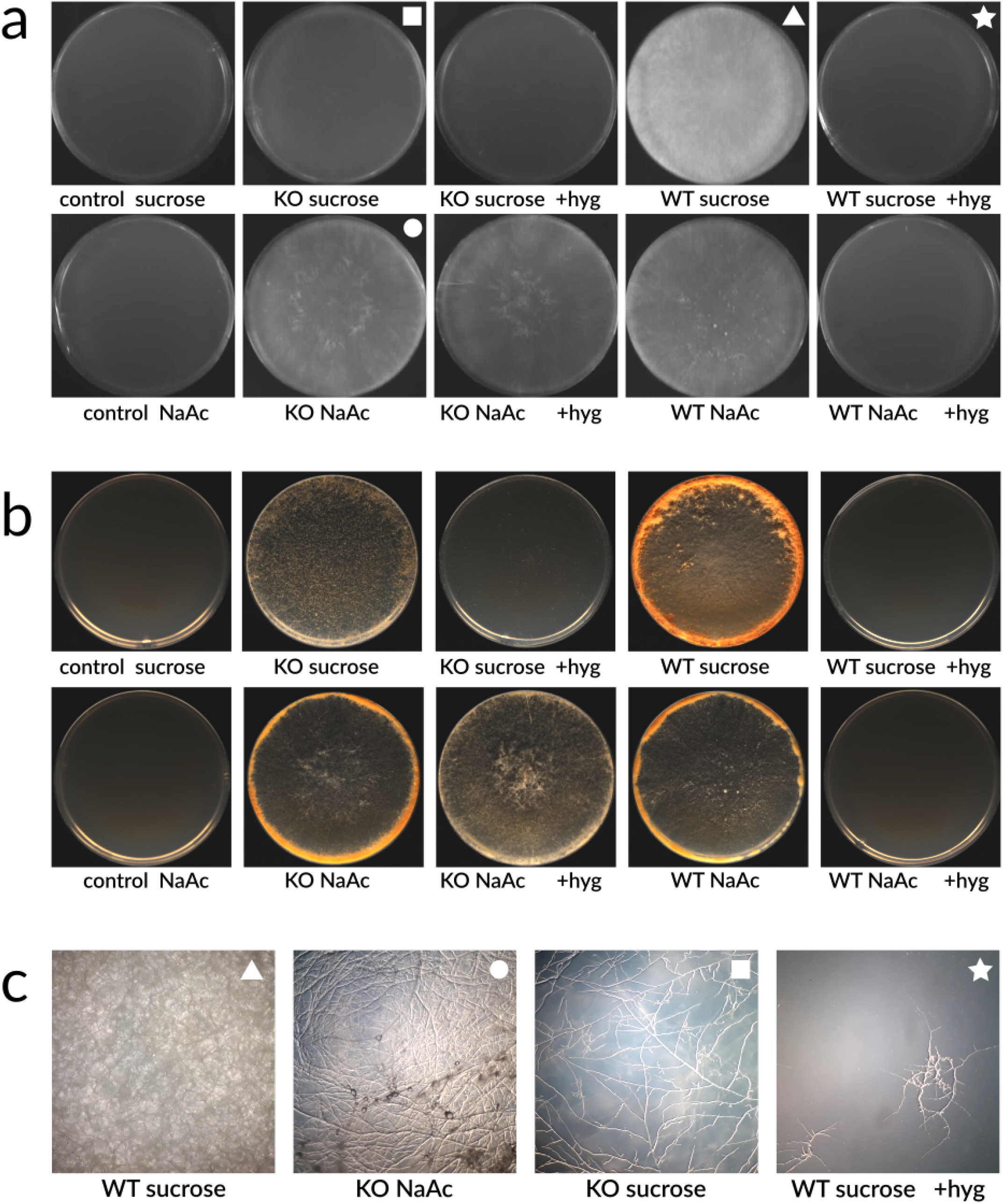
Differential growth assay results. **a**, During 72h in darkness, mycelia of *N. crassa* wildtype grows better on 1% sucrose than on 1% Na-Acetate (NaAc) (minimal Vogel’s media). Growth of a PX knockout is comparable to wildtype on NaAc, but is severely impeded on sucrose, indicating that PX is essential for pyruvate metabolism conveyed by the PDC. The knockout strain is engineered to be hygromycin-resistant, and supplementing with additional 300 *μ*M hygromycin is lethal to WT but has a minor effect on KO growth. **b**, Following an additional 72h in light, sporulation is also comparable in knockout and wildtype grown on NaAc. Sporulation is however distinctly different on sucrose. It appears that knockout of PX is detrimental but not lethal to *N. crassa*. **c**, Mycelial growth displays an indistinguishable phenotype despite the distinct differences in growth rate.

**Figure S9:**
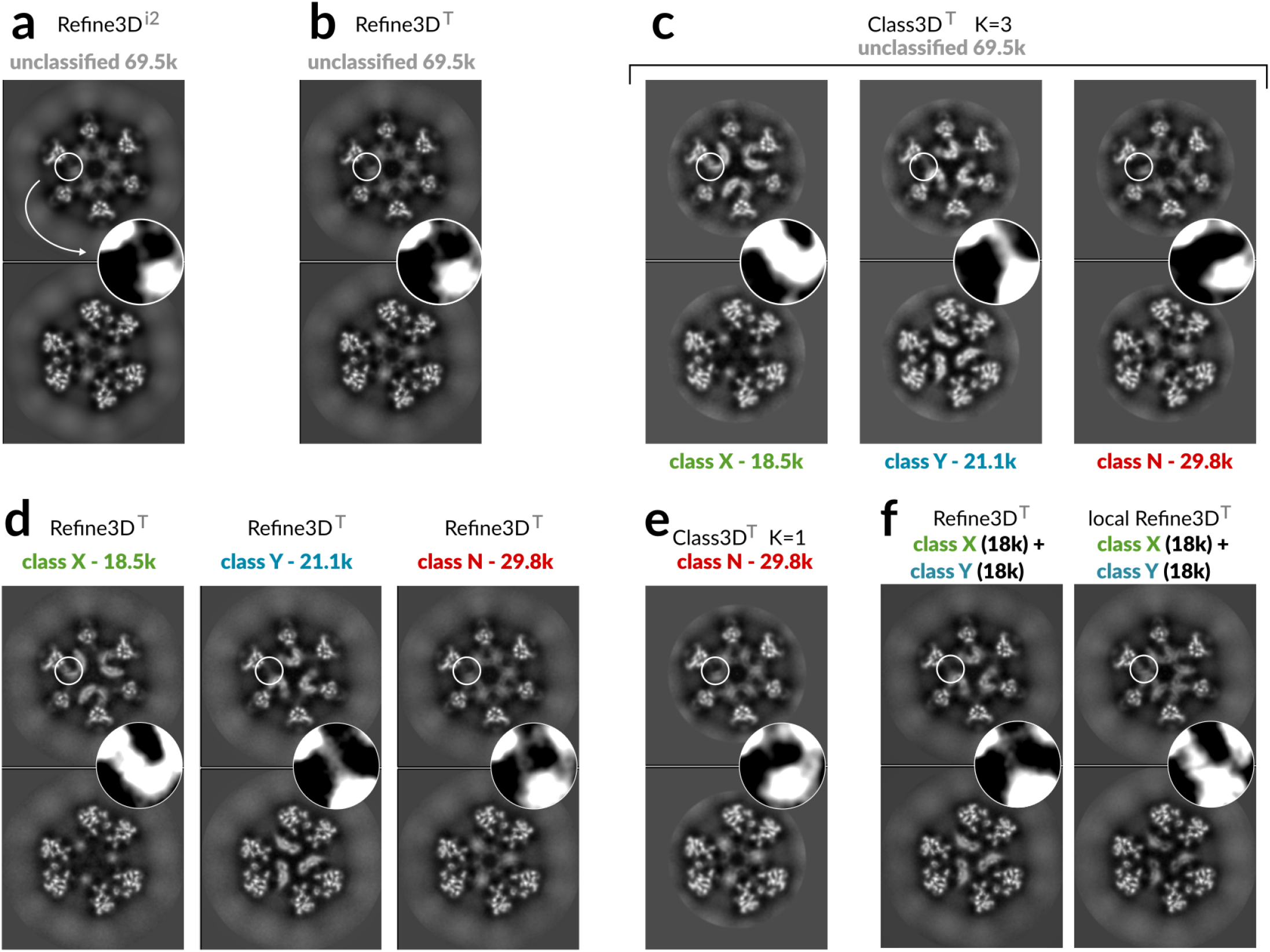
Reconstruction of recombined classes X and Y. **a**, Refinement of the endogenous PDC icosahedral ASU (I-ASU) shows over-symmetrization of a core-interior component. The inset shows a region of interest with increased contrast. **b**, Implicit symmetrization and non-tetrahedral particles cause the refinement of the tetrahedral ASU (T-ASU) to be similarly over-symmetrized prior to classification. **c**, Classification of the T-ASU is able to show distinct classes that do not superimpose (X and Y), and one yet over-symmetrized class (N). **d**, Refinement of the T-ASU from class N is similar but not identical to the classified reconstruction, indicating that some feature of the data in subclass N is hidden during classification. **e**, A classification of subclass N with a single class circumvents the use of local angular searches around a single orientation in the final steps of refinement, and does resemble the original classification. This then indicates that some data of subclass N is hidden during classification. **f**, Refinement of a re-combined dataset containing equal proportions of T-ASU classified X and Y (18k each), produce a Y-like reconstruction (compare inset to panel **d**), rather than a superimposed X/Y reconstruction. This shows efficient hiding of the X-classified ASU by weighted backprojection. By enforcing only local alignment to be used, data hiding by weighted back projection is prohibited, and the superimposed class X and Y is then also recovered in line with expectation. Note that no previous reconstruction shows a similar superposition. Local alignments in this case start from those found in individual X and Y refinements respectively.

## Notes

### Competing Interest Statement

The authors have declared no competing interest.

## References

1. Patel, K. P., O’brien, T. W., Subramony, S. H., Shuster, J. & Stacpoole, P. W. The spectrum of pyruvate dehydrogenase complex deficiency: clinical, biochemical and genetic features in 371 patients. Molecular genetics and metabolism 106, 385–394 (2012).

2. Gray, L. R., Tompkins, S. C. & Taylor, E. B. Regulation of pyruvate metabolism and human disease. Cellular and Molecular Life Sciences 71, 2577–2604 (2014).

3. Koukourakis, M. I. et al. Pyruvate dehydrogenase and pyruvate dehydrogenase kinase expression in non small cell lung cancer and tumor-associated stromal. Neoplasia 7, 1–6 (2005).

4. Tran, Q., Lee, H., Park, J. & Kim, S. H. Targeting cancer metabolism - revisiting the Warburg effects. Toxicological Research 32, 177–193 (2016).

5. Fisher-Wellman, K. H. et al. Mitochondrial glutathione depletion reveals a novel role for the pyruvate dehydrogenase complex as a key H2O2-emitting source under conditions of nutrient overload. Free Radical Biology and Medicine 65, 1201–1208 (2013).

6. Sutendra, G. & Michelakis, E. Pyruvate dehydrogenase kinase as a novel therapeutic target in oncology. Frontiers in Oncology 3, 1–11 (2013).

7. Kaplon, J. et al. A key role for mitochondrial gatekeeper pyruvate dehydrogenase in oncogene-induced senescence. Nature 498, 109–112 (2013).

8. Gu, Y. Q., Zhou, Z. H., McCarthy, D. B., Reed, L. J. & Stoops, J. K. 3D electron microscopy deposition and protein reveals the variable dynamics of the peripheral pyruvate dehydrogenase component about the core. Proceedings of the National Academy of Sciences of the United States of America 100, 7015–7020 (2003).

9. Milne, J. L. S. et al. Molecular architecture and mechanism of an icosahedral pyruvate dehydrogenase complex: a multifunctional catalytic machine. Embo Journal 21, 5587–5598 (2002).

10. Milne, J. L. S. et al. Molecular structure of a 9-MDa icosahedral pyruvate dehydrogenase subcomplex containing the E2 and E3 enzymes using cryoelectron microscopy. Journal of Biological Chemistry 281, 4364–4370 (2006).

11. Brautigam, C. A. et al. Structural insight into interactions between dihydrolipoamide dehydrogenase (E3) and E3 binding protein of human pyruvate dehydrogenase complex. Structure 14, 611–621 (2006).

12. Kato, M. et al. Structural basis for inactivation of the human pyruvate dehydrogenase complex by phosphorylation: role of disordered phosphorylation loops. Structure 16, 1849–1859 (2008).

13. Izard, T. et al. Principles of quasi-equivalence and Euclidean geometry govern the assembly of cubic and dodecahedral cores of pyruvate dehydrogenase complexes. Proceedings of the National Academy of Sciences of the United States of America 96, 1240–1245 (1999).

14. Jiang, J. S. et al. Atomic structure of the E2 inner core of human pyruvate dehydrogenase Complex. Biochemistry 57, 2325–2334 (2018).

15. Mattevi, A. et al. Atomic-structure of the cubic core of the pyruvate-dehydrogenase multienzyme complex. Science 255, 1544–1550 (1992).

16. Park, S. et al. Role of the Pyruvate Dehydrogenase Complex in Metabolic Remodeling: Differential Pyruvate Dehydrogenase Complex Functions in Metabolism. Diabetes & metabolism journal 42, 270–281 (2018).

17. Tareen, S. H. et al. Logical modelling reveals the PDC-PDK interaction as the regulatory switch driving metabolic flexibility at the cellular level. Genes & Nutrition 14, 27 (2019).

18. Demarcucci, O. & Lindsay, J. G. Component X -an immunologically distinct polypeptide associated with mammalian pyruvate-dehydrogenase multi-enzyme complex. European Journal of Biochemistry 149, 641–648 (1985).

19. Maeng, C. Y., Yazdi, M. A., Niu, X. D., Lee, H. Y. & Reed, L. J. Expression, purification, and characterization of the dihydrolipoamide dehydrogenase-binding protein of the pyruvate-dehydrogenase complex from saccharomyces cerevisiae. Biochemistry 33, 13801–13807 (1994).

20. Stoops, J. K. et al. On the unique structural organization of the Saccharomyces cerevisiae pyruvate dehydrogenase complex. Journal of Biological Chemistry 272, 5757–5764 (1997).

21. Brautigam, C. A., Wynn, R. M., Chuang, J. L. & Chuang, D. T. Subunit and catalytic component stoichiometries of an in vitro reconstituted human pyruvate dehydrogenase complex. Journal of Biological Chemistry 284, 13086–13098 (2009).

22. Hezaveh, S., Zeng, A. P. & Jandt, U. Full enzyme complex simulation: interactions in human pyruvate dehydrogenase complex. Journal of Chemical Information and Modeling 58, 362–369 (2018).

23. Hiromasa, Y., Fujisawa, T., Aso, Y. & Roche, T. E. Organization of the cores of the mammalian pyruvate dehydrogenase complex formed by E2 and E2 plus the E3-binding protein and their capacities to bind the E1 and E3 components. Journal of Biological Chemistry 279, 6921–6933 (2004).

24. Maeng, C. Y., Yazdi, M. A. & Reed, L. J. Stoichiometry of binding of mature and truncated forms of the dihydrolipoamide dehydrogenase-binding protein to the dihydrolipoamide acetyltransferase core of the pyruvate dehydrogenase complex from Saccharomyces cerevisiae. Biochemistry 35, 5879–5882 (1996).

25. Kato, M. et al. A synchronized substrate-gating mechanism revealed by cubic-core structure of the bovine branched-chain alpha-ketoacid dehydrogenase complex. Embo Journal 25, 5983–5994 (2006).

26. Vijayakrishnan, S. et al. Variation in the organization and subunit composition of the mammalian pyruvate dehydrogenase complex E2/E3BP core assembly. Biochemical Journal 437, 565–574 (2011).

27. Sheu, K. F. R., Hu, C. W. C. & Utter, M. F. Pyruvate-dehydrogenase complex activity in normal and deficient fibroblasts. Journal of Clinical Investigation 67, 1463–1471 (1981).

28. Randall, D. D., Rubin, P. M. & Fenko, M. Plant pyruvate-dehydrogenase complex purification, characterization and regulation by metabolites and phosphorylation. Biochimica Et Biophysica Acta 485, 336–349 (1977).

29. Neveling, U., Klasen, R., Bringer-Meyer, S. & Sahm, H. Purification of the pyruvate dehydrogenase multienzyme complex of Zymomonas mobilis and identification and sequence analysis of the corresponding genes. Journal of Bacteriology 180, 1540–1548 (1998).

30. Seifert, F. et al. Phosphorylation of serine 264 impedes active site accessibility in the E1 component of the human pyruvate dehydrogenase multienzyme complex. Biochemistry 46, 6277–6287 (2007).

31. Westergaard, M. & Mitchell, H. K. Neurospora-V - a synthetic medium favouring sexual reproduction. American Journal of Botany 34, 573–577 (1947).

32. Zheng, S. Q. et al. MotionCor2: anisotropic correction of beam-induced motion for improved cryo-electron microscopy. Nature Methods 14, 331–332 (2017).

33. Zhang, K. Gctf: Real-time CTF determination and correction. Journal of Structural Biology 193, 1–12 (2016).

34. Zivanov, J. et al. New tools for automated high-resolution cryo-EM structure determination in RELION-3. Elife 7 (2018).

35. Tegunov, D. & Cramer, P. Real-time cryo-EM data pre-processing with Warp. Nature Methods 16, 1146–1152 (2019).

36. Emsley, P., Lohkamp, B., Scott, W. G. & Cowtan, K. Features and development of Coot. Acta Crystallographica Section D-Biological Crystallography 66, 486–501 (2010).

37. Adams, P. D. et al. PHENIX: a comprehensive Python-based system for macromolecular structure solution. Acta Crystallographica Section D-Biological Crystallography 66, 213–221 (2010).

38. Huang, C. C. et al. UCSF Chimera. Abstracts of Papers of the American Chemical Society 238 (2009).

39. Goddard, T. D. et al. UCSF ChimeraX: meeting modern challenges in visualization and analysis. Protein Science 27, 14–25 (2018).

40. El-Gebali, S. et al. The Pfam protein families database in 2019. Nucleic Acids Research 47, D427–D432 (2019).

41. Waterhouse, A. M., Procter, J. B., Martin, D. M. A., Clamp, M. & Barton, G. J. Jalview version 2-a multiple sequence alignment editor and analysis workbench. Bioinformatics 25, 1189–1191 (2009).

42. Ma, J. Z., Wang, S., Wang, Z. Y. & Xu, J. B. Protein contact prediction by integrating joint evolutionary coupling analysis and supervised learning. Bioinformatics 31, 3506–3513 (2015).

43. Hu, M. X. et al. A particle-filter framework for robust cryo-EM 3D reconstruction. Nature Methods 15, 1083–1089 (2018).

